# Rrp6p/Rrp47p constitutes an independent nuclear turnover system of mature small non-coding RNAs in *Saccharomyces cerevisiae*

**DOI:** 10.1101/2020.12.13.422512

**Authors:** Anusha Chaudhuri, Subhadeep Das, Mayukh Banerjea, Biswadip Das

## Abstract

In *Saccharomyces cerevisiae,* the nuclear exosome/Rrp6p/TRAMP participates in the 3’-end processing of several precursor non-coding RNAs. Here we demonstrate that the depletion of nucleus-specific 3’→5’ exoribonuclease Rrp6p and its cofactor, Rrp47p led to the specific and selective enhancement of steady-state levels of mature small non-coding RNAs (sncRNAs) that include 5S and 5.8S rRNAs, snRNAs and snoRNAs, but not 18S and 25S rRNAs. Most importantly, their steady-state enhancement does not require the exosome, TRAMP, CTEXT, or Rrp6p-associated Mpp6p. Rrp6p/47p-dependent enhancement of the steady-state levels of sncRNAs is associated with the diminution of their nuclear decay-rate and requires their polyadenylation before targeting by Rrp6p, which is catalyzed by both the canonical and non-canonical poly(A) polymerases, Pap1p and Trf4p. Consistent with this finding, we also demonstrated that Rrp6p and Rrp47p exist as an exosome-independent complex. Thus, Rrp6p-Rrp47p defines a core nuclear exosome-independent novel turnover system that targets the small non-coding RNAs.

## Introduction

In baker’s yeast *Saccharomyces cerevisiae*, rDNA locus consists of 100 to 200 copies of the 9.1 kb long rDNA unit containing the 35S rRNA operon and two non-transcribed spacer units (NTSs), which are separated by 5S rRNA gene. The 35S operon consists of 18S, 5.8S, and 25S rRNA genes (1–3). Its transcription by RNA Polymerase I produces a 35S primary transcript, which undergoes extensive endonucleolytic cleavage and exonucleolytic processing reactions (3–7) to yield various mature rRNA species (1, 2, 15–19, 3, 8–14). In contrast to the 35S operon, the 5S rRNA gene is transcribed in the opposite direction by RNA polymerase III as a precursor, extended by ten nucleotides at its 3’-end and requires the product of the *RNA82* gene for their removal (20) followed by several modification events.

The mature 3’-ends of individually transcribed snoRNAs are coupled to their transcription termination (21–26) mediated by Nrd1p-Nab3p-Sen1p (NNS) complex in association with a few cleavage polyadenylation components (26). The dissociation of the RNA from the DNA promoted by Sen1p (27) leads to further recruitment of Pcf11p, the nuclear exosome, TRAMP, and Rrp6p to their maturing 3’-terminus (28–31) resulting in its trimming until the snoRNA secondary structure (32). 3’-end maturation of the intronic snoRNAs, in contrast, is initiated by the debranching of the intron lariat by Dbr1p and Rnt1p (33–35) and snoRNAs release followed by their exonucleolytic processing at the 3’-end. NNS complex subsequently binds to the 3’-end of these debranched RNAs to stimulate their 3’-end processing (36) although the involvement of the exosome/Rrp6p in this process is uncertain (21). All the snRNA genes (U1, U2, U4, and U5), except U6 snRNA, are transcribed by RNA Pol II, whereas the latter is transcribed by RNA Pol III (37, 38). The formation of the 3’-ends of the snRNA primary transcripts are mediated co-transcriptionally by the NNS complex that eventually leads to the formation of the precursor snRNAs with a long 3’-extension, which are subsequently processed by the exosome and Rrp6p (26).

The nuclear exosome, implicated in the 3’-end processing events of many of these non-coding RNA precursors and processing intermediates, consists of nine core subunits arranged in two layers stacked-donuts with a common central channel. Three cap subunits, Rrp40p, Rrp4p, and Csl4p, are placed as a trimeric cap on the top of the remaining six subunits, Rrp41p, Rrp42p, Mtr3p, Rrp43p, Rrp46p, and Rrp45p, which collectively form the bottom hexameric ring structure (39). Strikingly, the core exosome (Dubbed Exo-9) lacks the catalytic activity for RNA hydrolysis despite the presence of the putative active site for 3’→5’ exoribonuclease in each subunit (39, 40). The entire exosome complex’s catalytic activity resides in the tenth and eleventh subunits, Dis3p/Rrp44p and Rrp6p, respectively, each of which makes contact with the Exo-9 structure from opposite sides (40, 41). While Dis3p/Rrp44p is associated with both nuclear and cytoplasmic forms of the exosome, Rrp6p is specifically associated with the nuclear form of the RNA exosome (15, 42). Additionally, exosome is associated with several ancillary factors, Lrp1p/Rrp47p (specific to nuclear exosome), Mpp6p (specific to nuclear exosome), and *SKI* complex (specific to cytoplasmic exosome) (43). Catalytic specificity of the exosome is brought about by ancillary protein complexes, all of which target a distinct set of RNA substrates to the exosome. Two such complexes found in the nucleus of *S. cerevisiae* are the TRAMP complex and the CTEXT complex. TRAMP (**<u>TR</u>**f4p/5p-**<u>A</u>**ir1p/2p-**<u>M</u>**tr4p-**<u>P</u>**olyadenylation) complex consisting of non-canonical poly(A) polymerase, Trf4p/Trf5p, a zinc knuckle protein Air1p/Air2p, and the RNA helicase Mtr4p (19, 30, 44, 45) assists the exosome to target faulty mRNA transcripts yielded at the earlier phase of mRNP biogenesis (46) and in the processing/modification events of other non-coding RNAs. CTEXT (**<u>C</u>**bc1p-**<u>T</u>**if4631p-dependent **<u>EX</u>**osomal **<u>T</u>**argeting) (previously termed as DRN) (46), in contrast, consists of nuclear cap-binding protein Cbc1p/2p (47, 48), shuttling proteins Tif4631p/ Upf3p (49), a DEAD-box RNA helicase, Dbp2p (50) and meiotic protein Red1p (51), which collectively participate in the degradation of aberrant mRNAs produced at the late phase of mRNP biogenesis (46).

The “Core exosome model,” subsisting for a long time, stipulated the obligatory and collective contribution of each of the eleven exosome subunits for the structural integrity and catalytic activity of this complex in all RNA metabolic functions (15, 18, 59–64, 41, 52–58). However, a large body of experimental data substantiates collective inferences that were not predicted and anticipated by the ‘core exosome model’ (30, 44, 72, 64–71). All of this evidence instead strongly support the idea that specific subunits of the core exosome may physically exist either as a monomer or as a separate complex in association with one or more exosomal or non-exosomal components (39, 73–78). Furthermore, many exosome components function independently of the core exosome by targeting distinct sets of RNA substrates (67, 68, 71, 78–80), which are not targeted by the entire exosome complex. Moreover, the intracellular localization profiles and copy numbers of the various exosomal subunits vary, indicating that individual subunits may have Exo-11 complex-independent function (63, 81–83). Collectively, all these findings led to an alternative ‘exozyme’ hypothesis that demands that in addition to its role as a part of the exosome complex, some exosome components assemble into and function as independent complexes (65). In good agreement with the ‘exozyme’ concept, the nuclear exosome component Rrp6p alone was shown to independently process the 3’-end of the 5.8S+30 RNA intermediate (18) and degrade the polyadenylated version of ribosomal RNAs (84). However, whether the Rrp6p-dependent decay of polyadenylated rRNA represents an independent function from core exosome was not addressed and known (84). A later study indeed revealed that some of the degradation activities of Rrp6p occur independently of core exosome (68), which was supported by the finding that cells depleted of Rrp6p accumulate poly(A)^+^ rRNA degradation intermediates, which are different from those found in cells lacking either Dis3p or the core exosome component Rrp43p (68).

Furthermore, depletion of Rrp44p in the absence of Rrp6p led to the synergistic enhancement of the steady-state levels of decay intermediates common to core exosome and Rrp6p but did not have any effect on the levels of decay-intermediates, which are Rrp6p specific. Strikingly, disrupting the physical interaction between Rrp6p and core exosome did not affect the processing of the 3’-end of 5.8S rRNA and snoRNAs and some of these Rrp6p-specific decay intermediates. Collectively, these findings affirmed that Rrp6p carry out these vital activities independently of core exosome (68).

In this study, we address if Rrp6p carries out the degradation of the mature and precursor non-coding RNA species (rRNAs, snRNAs, and snoRNAs) independently of the core-exosome by systematically analyzing the functional contributions of the Rrp6p, and components of core exosome, TRAMP, and CTEXT in the steady-state accumulation of rRNAs, sn- and snoRNAs. Our investigation revealed a dramatic accumulation of the polyadenylated forms of only small non-coding stable RNAs (5S and 5.8S rRNAs, snRNAs and snoRNAs) only in the *rrp6-*Δ and *rrp47-*Δ strains but not in the strains carrying the mutant alleles of core exosome, TRAMP, CTEXT, and Rrp6p-associated Mpp6p. This finding suggests that Rrp6/47p coordinates the degradation of the polyadenylated version of the mature form of only the small non-coding RNAs independent of the core exosome. Interestingly, thorough analyses of the global transcriptome from a previously published RNA-seq dataset revealed a dramatic accumulation of sequence reads mapped to the mature regions of some of these snoRNAs in *rrp6*-Δ yeast strains. Polyadenylation of the targeted small ncRNA species is crucial, which requires the involvement of both the canonical and non-canonical polyA-polymerase, Pap1p and Trf4p. We believe that Rrp6/47p constitutes a novel nuclear turnover system of small non-coding RNAs that target the fraction of the small non-coding RNAs that fail to undergo further nuclear maturation.

## Results

### Steady-state levels of 5S, 5.8S rRNAs, snRNAs, and select snoRNAs display a dramatic enhancement in *rrp6-*Δ strain

An unanticipated and dramatic enhancement of steady-state levels of 5S/5.8S rRNA exclusively in an *rrp6-*Δ yeast strain in our laboratory inspired this investigation. This unexpected observation was consistently noted during traditional mRNA decay experiments, in which these small RNA species, being the transcripts of either RNA Pol III or RNA Pol I, were used as an internal control. Although initially this finding was considered as an artifact, this observation proved very reproducible in diverse genetic strain backgrounds and was reminiscent of the Rrp6p-dependent degradation of the polyadenylated form of rRNAs reported previously (84). This observation prompted us to hypothesize that the steady-state increment of these small non-coding RNAs in the *rrp6-*Δ strain represents their degradation in the nucleus that requires Rrp6p. To test our hypothesis, we determined the steady-state levels of all the ribosomal RNAs in the wild-type strain and the strains carrying the mutations in the components of the core exosome (*rrp6*-Δ, *rrp4-1*, and *GAL10::RRP41*), TRAMP (*mtr4-1/dob1-1*, *trf4-*Δ, *trf5-*Δ, and *air1*-Δ), CTEXT (*cbc1*-Δ and *tif4631*-Δ) by qRT-PCR using cDNA prepared from these strains using either random hexanucleotide primer or an oligo-dT anchor primer (Fig. 1A) followed by the quantitative amplification of the amplicons corresponding to the mature sequences of each of the ribosomal RNAs (Fig. 1B). Therefore, these assays enable us to precisely quantify the populations of both the total and polyadenylated forms of the mature and precursors rRNA species in various strains. As shown in Figure 1C and D, the steady-state levels of neither total nor the polyadenylated version of 18S and 25S rRNAs showed any significant level of alteration in any of these strains. The steady-state levels of both total (polyadenylated and non-poly(a) fraction together) and polyadenylated versions of mature and precursor 5S and 5.8S rRNAs, in contrast, displayed a dramatic enhancement in the strains depleted of Rrp6p (*rrp6*-Δ strain) (~7 fold from cDNA prepared using the random primer and ~175 fold from cDNA prepared using oligo-dT primer). The level of augmentation, however, is relatively moderate (~2 and ~1.8 folds strains from cDNA prepared using the random primer and ~7 and ~2 folds in cDNA prepared using oligo-dT primer) in the strains carrying *rrp4-1* and *GAL10::RRP41* strains, respectively. Interestingly, the steady-state levels of the 5S and 5.8S rRNAs were found to exhibit a very mild enhancement in *trf4-*Δ strain, which does not appear evident in the histogram scale presented in Fig. 1 (see Fig. 8 and below). Failure of the *upf1*-Δ to enhance the levels of these two small ribosomal RNAs implied that the decay is not cytoplasmic (Fig. 1C and D).

**Figure 1:**
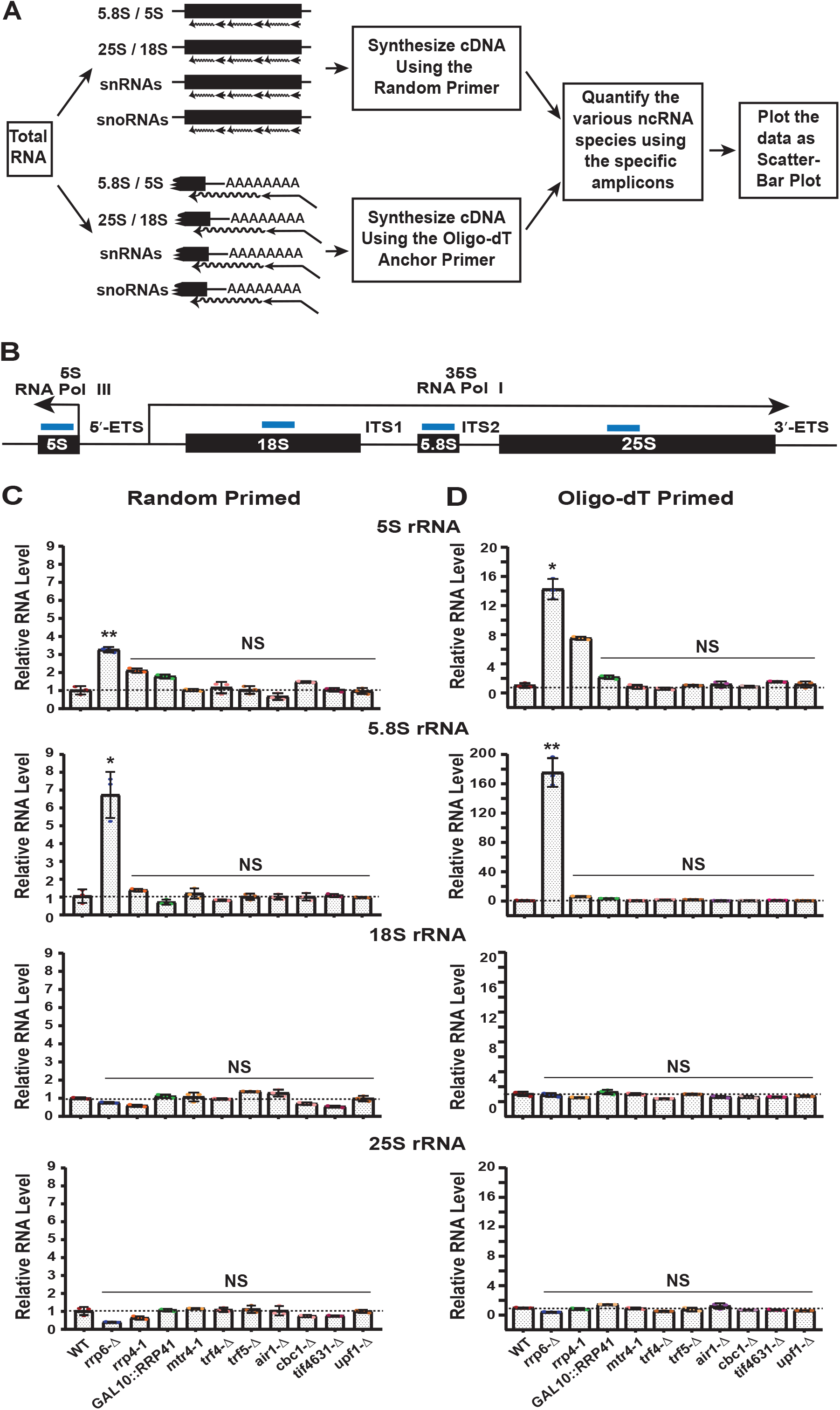
Polyadenylated version of mature forms of low molecular weight ribosomal RNAs accumulate in an *rrp6*-Δ yeast strain. (A) Schematic diagram showing the general outline of the experimental approach undertaken in the present work to detect the abundance of the various non-coding RNA species in WT and different yeast strains. (B) Schematic representation of the genes for the 5S, 18S, 5.8S, and 25S rRNAs depicting the amplicons’ locations (thick blue lines on top of each of these genes) used to determine the abundances of the mature form of those RNAs by qRT-PCR. Scattered/Bar plot revealing the steady-state levels of the mature form of various rRNAs estimated from 2 ng cDNA samples prepared using random hexanucleotide primers (C) or oligo-dT_30_ anchor primer (D) by qRT-PCR using the amplicons shown in Fig.1B from the wild type strain and strains carrying mutations in the component of the nuclear exosome (*rrp6*-Δ, *rrp4-1*, *GAL10::RRP41*), TRAMP complex (*mtr4-1*, *trf4-*Δ, *trf5-*Δ, and *air1*-Δ) and CTEXT (*cbc1*-Δ, *tif4631*-Δ). *SCR1* (in the case of Random Primer) and *ACT1* mRNA (in the case of Oligo dT Primer) were used as the internal control. The abundance of these ncRNAs in *upf1-*Δ yeast strain was used as a negative control. Normalized values of each of the ncRNAs in the wild-type yeast strain were set to 1. Three independent cDNA preparations (biological replicates, n = 3 and in few cases 4) were used to determine the levels of various ncRNAs. The statistical significance of difference reflected in the ranges of P values estimated from Student’s two-tailed t-tests for a given pair of test strains for each message are presented with the following symbols, *<0.05, **<0.005, and ***<0.001; NS, not significant.

Next, we addressed if the steady-state levels of other non-coding RNAs are also affected by Rrp6p dependent manner. We determined the steady-state levels of total and polyadenylated cellular populations (both mature and precursors) of different snRNAs, U1, U2, U4, U5 (sm class) and U6 (Lsm class) and two different categories of snoRNAs, snR10 (representatives of H/ACA box snoRNAs) and snR13 (a representative of C/D box snoRNA) from the cDNA prepared using both the random and oligo-dT12-18 anchor-primer from various strains as indicated in Figure 1A. As shown in Fig. 2A-B, total populations of all the mature sn- and snoRNAs exhibited dramatic levels of enhancement (~4 to 18 folds for snRNAs and ~6 to 10 folds for snoRNAs estimated using the random primer and ~25 to 110 folds for snRNAs and ~80 to 90 folds for snoRNAs as estimated from oligo-dT primer) in their abundance in strains carrying the *rrp6*-Δ allele. Mutations in no other genes displayed any enhancement of any total populations of sn- or snoRNAs (Fig. 2A-B) except a mild increment found in *trf4-*Δ strain (see Fig. 8 and below). Thus, it appears that the steady-state levels of the total cellular populations of small non-coding RNAs are dramatically increased in a yeast strain carrying the *rrp6-*Δ allele. This finding led us to interpret that Rrp6p constitutes an exosome-independent nuclear degradation system that targets explicitly small non-coding RNAs like 5S, 5.8S rRNAs, snRNAs, and snoRNAs. The other exosomal components (except Rrp4, which target mature 5S rRNA) and cofactors do not participate in this degradation. Thus our experimental findings indicate a nuclear Rrp6p-dependent degradation system that acts on the polyadenylated forms of the low-molecular-weight non-coding RNAs.

**Figure 2:**
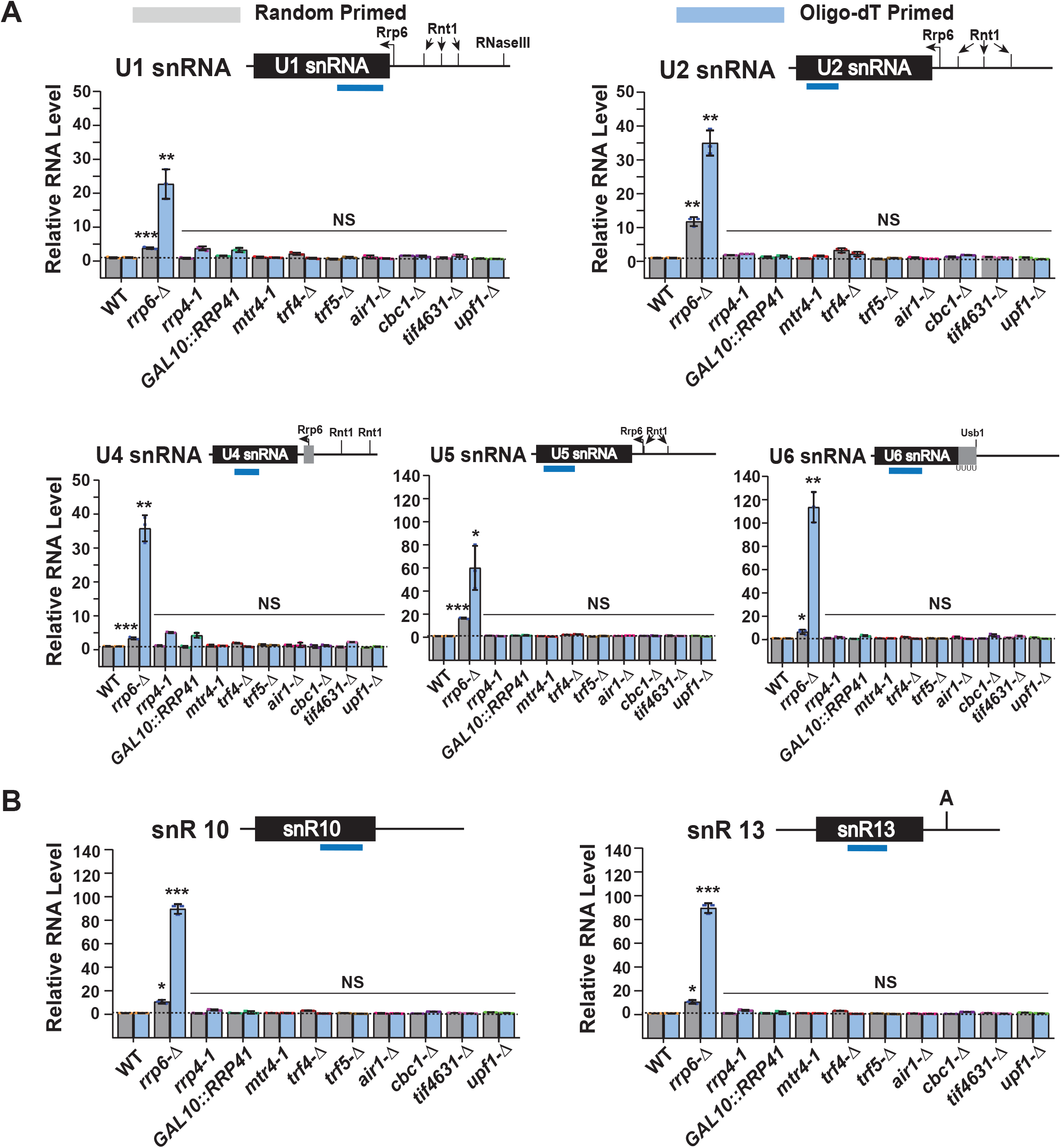
Polyadenylated version of mature forms various snRNAs and snoRNAs accumulate in an *rrp6*-Δ yeast strain. Scattered/Bar plot revealing the steady-state levels of the mature form of various (A) snRNAs and (B) snoRNAs estimated from the 2 ng cDNA samples prepared using either random hexanucleotide primers (grey bars) or oligo-dT_30_ anchor primer (light blue bars) by qRT-PCR using the amplicons shown in the schematic figure presented on top of each graph from indicated yeast strains. The solid black box downstream of each ncRNA gene indicates the Rrp6p processing site. *SCR1* (in the case of Random Primer) and *ACT1* mRNA (in the case of Oligo dT Primer) were used as the internal control. The abundance of these ncRNAs in *upf1-*Δ yeast strain was used as a negative control. Normalized values of each of the ncRNAs in the wild-type yeast strain were set to 1. Normalized values of each of the ncRNAs in the wild-type yeast strain were set to 1. Three independent cDNA preparations (biological replicates, n = 3 and in some cases 4) were used to determine the levels of various ncRNAs. The statistical significance of difference reflected in the ranges of P values are presented with the following symbols, *<0.05, **<0.005, and ***<0.001; NS, not significant.

### Rrp6p targets the mature forms of the small non-coding RNAs

To assess if the steady-state enhancement of the precursor/mature population of small non-coding RNAs in the *rrp6*-Δ strain is a direct consequence of accumulating their 3’-extended precursors or their mature forms, we precisely determined the steady-state levels of the precursors. Therefore we determined the steady-state levels of their precursors from cDNA samples from various strains primed with either the random or oligo-dT primers as before, followed by qRT-PCR. However, in these assays, we used specific primer sets those span the mature-precursor regions of each RNA species (indicated on top of each histogram in Figs. 3 and 4 as thick blue lines below the RNA sequence). For 5.8S rRNA, two different precursors species were evaluated; one was corresponding to 7SS/L precursor/intermediate (dubbed pre-5.8S RNA-II) (Fig. 3A). The other one was 5.8S+30 intermediate (dubbed pre-5.8S RNA-I), which was previously shown to be exonucleolytically processed by Rrp6p (84, 85) (Fig. 3A).

**Figure 3:**
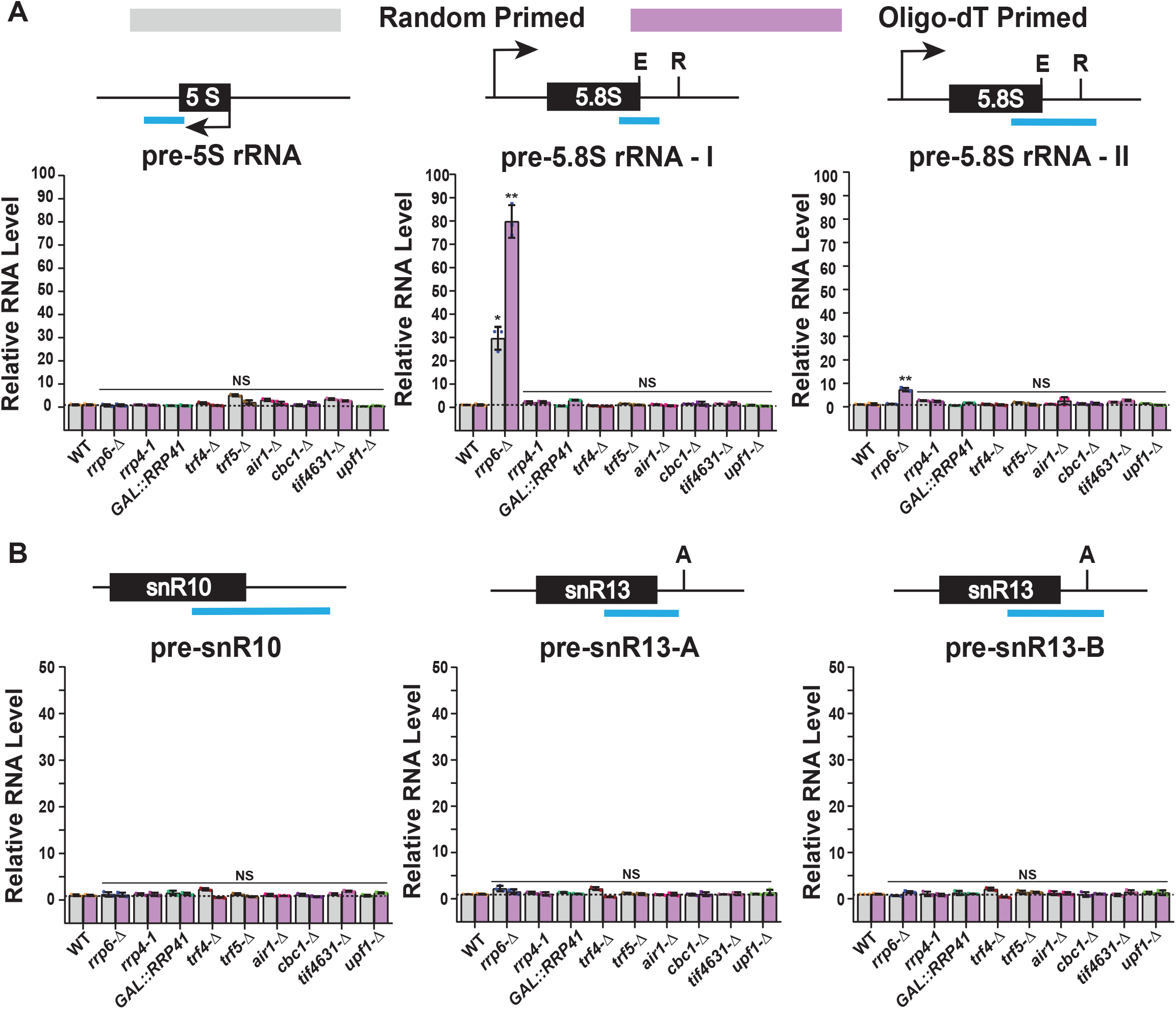
Steady-state levels of 3’-extended precursor forms of low molecular weight rRNAs and different test snoRNAs in various yeast strains. Scattered/Bar plot revealing the steady-state levels of 3’-extended precursor forms of (A) 5S and 5.8S small rRNAs and (B) two snoRNAs in indicated yeast strains. The RNA levels were estimated from the 2 ng cDNA samples prepared using random hexanucleotide primers (grey bars) or oligo-dT_30_ anchor primer (pink bars). Various amplicons used in subsequent qRT-PCR reactions covering partly the mature region and partly the 3’-extended precursor regions of each RNA species are shown in the schematic figures presented on top of each graph. E and R indicate the mature 3’-termini and Rrp6 processing sites of 5S and 5.8S rRNA genes, respectively. A indicates 3’-processing site of snR13 gene. *SCR1* (in the case of Random Primer) and *ACT1* mRNA (in the case of Oligo dT Primer) were used as the internal control. The abundance of these ncRNAs in *upf1-*Δ yeast strain was used as a negative control. Normalized values of each of the ncRNAs in the wild-type yeast strain were set to 1. Normalized values of each of the ncRNAs in the wild-type yeast strain were set to 1. Three independent cDNA preparations (biological replicates, n = 3) were used to determine the levels of various ncRNAs. The statistical significance of difference reflected in the ranges of P values are presented with the following symbols, *<0.05, **<0.005, and ***<0.001; NS, not significant.

**Figure 4:**
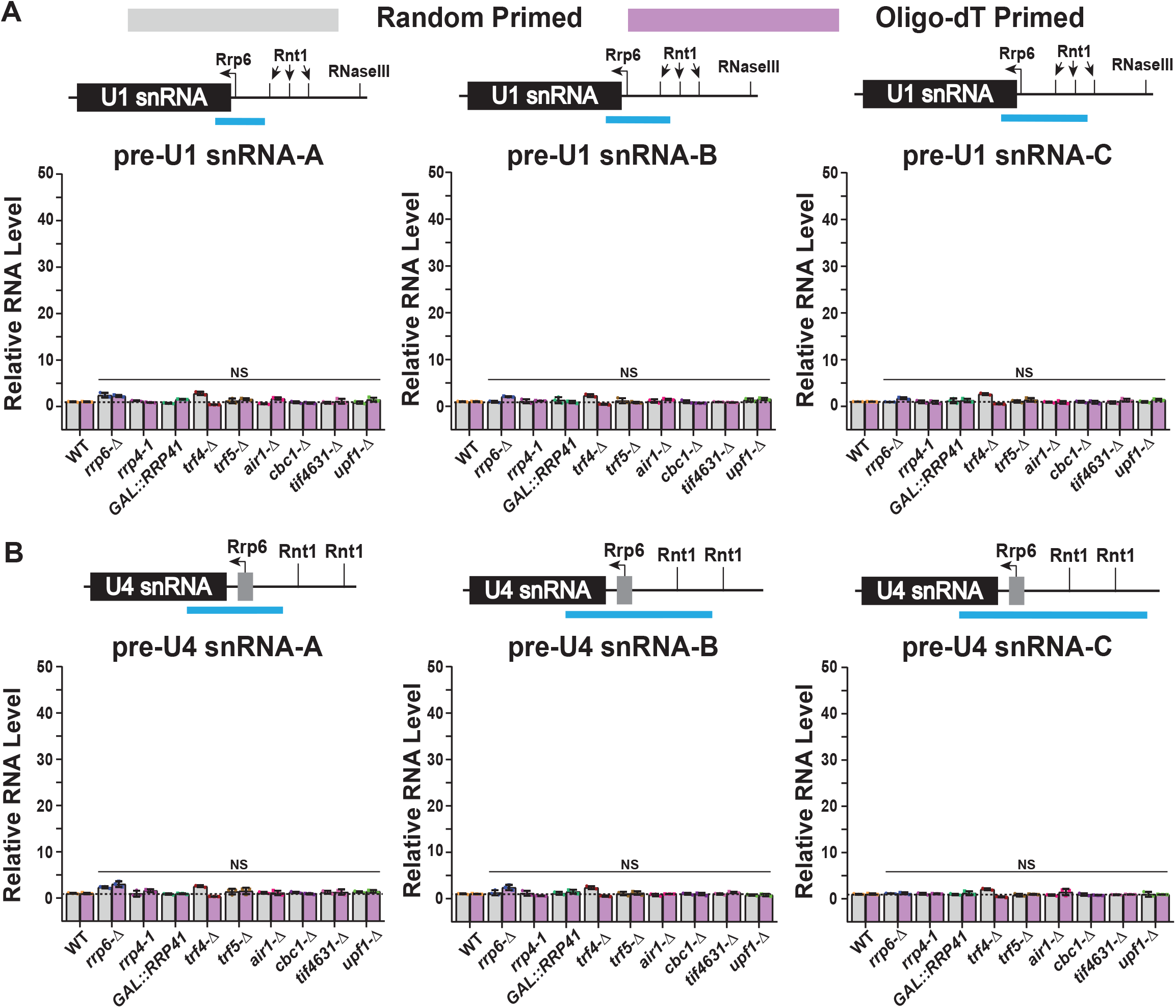
Steady-state levels of 3’-extended precursor forms of different test snRNAs in various yeast strains. Scattered/Bar plot revealing the steady-state levels of 3’-extended precursor forms of (A) snRNA U1 and (B) snRNA U4 in the indicated yeast strains. The RNA levels were estimated from the 2 ng cDNA samples prepared using random hexanucleotide primers (grey bars) or oligo-dT_30_ anchor primer (pink bars). Various amplicons used in subsequent qRT-PCR reactions covering partly the mature region and partly the 3’-extended precursor regions of each RNA species are shown in the schematic figures presented on top of each graph. Different 3’-end processing sites indicating cleavage by Rrp6p, Rnt1p, and RNaseIII are denoted in the respective schematic figures. *SCR1* (in the case of Random Primer) and *ACT1* mRNA (in the case of Oligo dT Primer) were used as the internal control. The abundance of these ncRNAs in *upf1-*Δ yeast strain was used as a negative control. Normalized values of each of the ncRNAs in the wild-type yeast strain were set to 1. Three independent cDNA preparations (biological replicates, n=3) were used to determine the levels of various ncRNAs. The statistical significance of difference reflected in the ranges of P values are presented with the following symbols, *<0.05, **<0.005, and ***<0.001; NS, not significant.

Our findings revealed that the steady-state levels of total and polyadenylated pre-5S and pre5.8S-II (presumably corresponding to 7SS/L intermediate) remained unaltered in all mutant yeast strains carrying the mutations in components of the core exosome, TRAMP, and CTEXT, including the *rrp6*-Δ strain. In contrast, a ≈30 and ≈80 fold increment in the level of pre-5.8S rRNA-I was noted in *rrp6*-Δ strain using the random-primed and oligo-dT primed cDNA, respectively. This data indicates that although a substantial amount of pre5.8S rRNA-I occurs in the *rrp6*-Δ strain due to an accumulation of the 5.8S+30 intermediate RNA species as reported before (84, 85), no significant accumulation of pre-5.8S rRNA-II species was noted. This finding thus prompted us to precisely evaluate the levels of the precursor species of different select sno- and snRNAs using a similar strategy using amplicons spanning from mature regions to the extended precursor regions (indicated on top of each histogram in Figs. 3B, 4 and supplementary Fig. S1). Our analysis showed that none of these precursor RNA species’ steady-state levels were increased in the *rrp6*-Δ yeast strain.

Furthermore, a careful comparison of the enhancement of total 5.8S RNA species (~ 200 fold as shown in Fig. 2C-D) with that of the pre-5.8S rRNA-II species (~ 30-80 folds as shown in Fig. 3A) indicates that a significant amount of mature 5.8S/5S/sn-/snoRNA species accumulate in *rrp6-*Δ strain. This finding warrants more careful analysis/ comparison of the steady-state levels of different amplicons corresponding to various regions spanning the mature sequence of diverse ncRNA species with different amplicons corresponding to various lengths encompassing the 3’-extended precursor regions (Fig. 5). Thus, using these amplicons, we precisely determined the levels of the mature and 3’-extended precursor forms of these RNAs in WT and *rrp6-*Δ strains using only oligo-dT primed cDNA and presented them as the normalized RNA species accumulated in the *rrp6*-Δ strain relative to WT (expressing their abundance in WT as 1) (Fig. 5). The normalized value of each of the RNA species was plotted as log2 fold change to accommodate a wide range of abundance values for diverse ncRNAs. For 5S rRNA species, no accumulation of any 3’-extended pre-5S RNA in comparison to mature 5S RNA was noted in *rrp6-*Δ yeast strain, thus clearly suggesting that a significant amount of mature 5S rRNA species was found to accumulate in the strain deficient of Rrp6p (Fig. 5A). In the case of 5.8S rRNA, we tested and compared the abundances of two different amplicons in the mature regions (Mature A and B) with three different regions of 3’-extended precursor forms (Pre-5.8S A, B, and C). As expected from the previous measurement as described above, the levels of Mature 5.8S A and B displayed 60 to 130 fold accumulation; the levels of the Pre-5.8S A, B, and C revealed 30, 3, and 1.7 fold accumulation in the *rrp6*-Δ strain. Together, these data strongly indicate a dramatic accumulation of the mature polyadenylated species of both 5S and 5.8S rRNA in the *rrp6*-Δ strain that Rrp6p targets and degrades the mature forms of both of these small rRNA species. The accumulation of pre-5.8S A amplicons (corresponding to the 5.8S+30 intermediate) in *rrp6*-Δ strain was associated with the exosome-independent Rrp6p-dependent processing/trimming of last 3’-30 nucleotides of pre-5.8S rRNA species (84–86).

**Figure 5:**
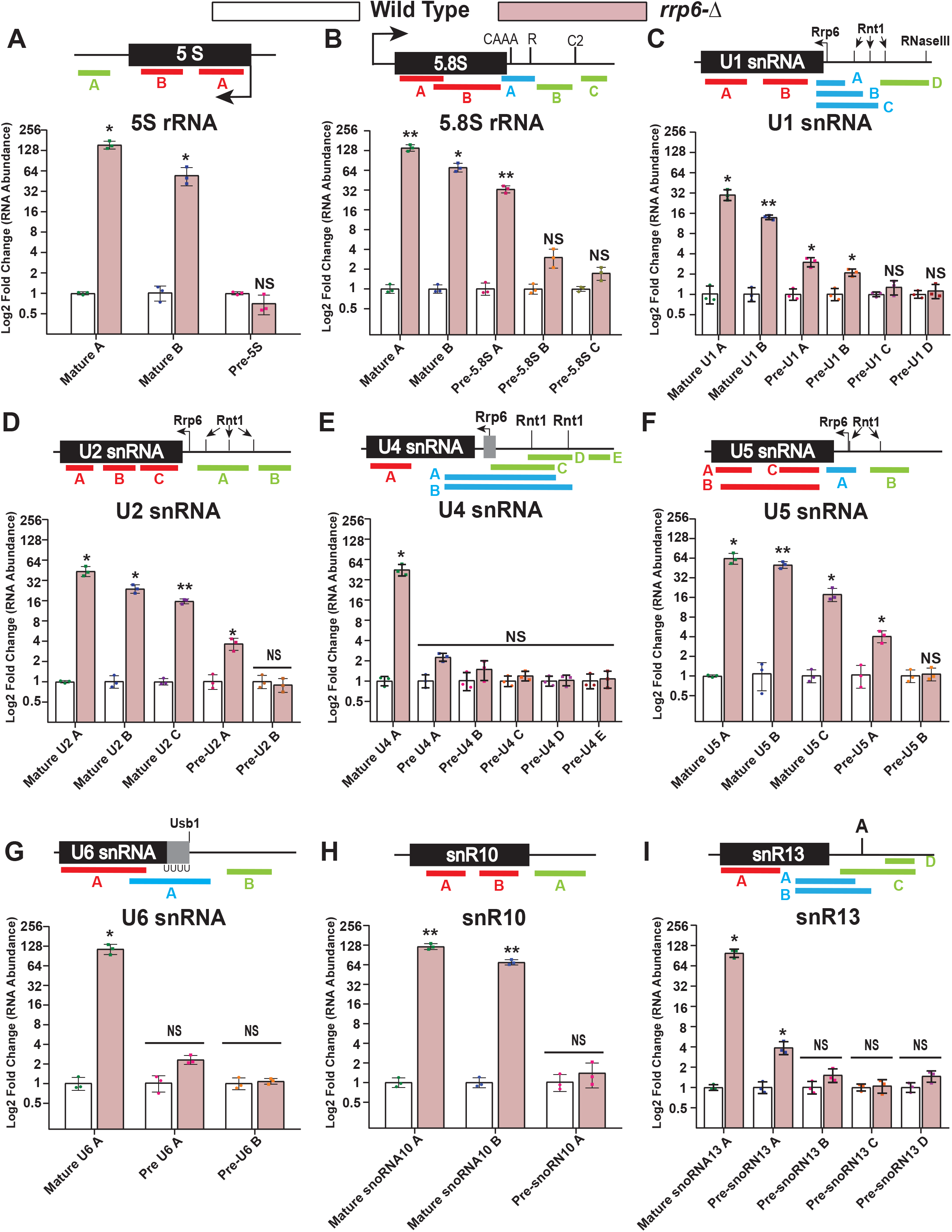
Comparison of steady-state levels of polyadenylated mature forms with that of various 3’-extended precursor forms of low molecular weight ncRNAs in *rrp6-*Δ yeast strains. Scattered/Bar plot revealing the steady-state levels of amplicons corresponding to the various regions in mature and 3’-extended precursor species of various small ncRNAs in WT (white bars) and *rrp6-*Δ (magenta bars) yeast strains. The RNA levels were estimated from the 2 ng cDNA samples prepared using the oligo-dT_30_ anchor primer. Various amplicons used in subsequent qRT-PCR reactions are shown in the schematic figures presented on top of each graph (red indicates mature amplicons, blue indicates intermediate amplicons covering partly the mature region and partly the 3’-extended precursor regions, and green indicates extended 3’-end precursor amplicons). Different 3’-end processing sites showing processing site of Rrp6p and cleavage sites of Rnt1p, RNase III, Usb1p, and others are denoted in the respective schematic figures. *ACT1* was used as internal control. Normalized values of each of the ncRNAs in the wild-type yeast strain were set to 1. Three independent cDNA preparations (biological replicates, n = 3) were used to determine the levels of various ncRNAs. The statistical significance of difference reflected in the ranges of P values are presented with the following symbols, *<0.05, **<0.005, and ***<0.001; NS, not significant.

Importantly, all sn- and snoRNAs also displayed a similar and dramatic accumulation of exclusively mature RNA species in the *rrp6*-Δ strain relative to their levels in WT strain (the abundance of their polyadenylated mature forms varied from 20 to 130 fold for different amplicons encompassing different snRNAs) (Fig. 5C-I). In every case, the proximal (to the mature 3’-end) 3’-extended precursor species of these RNAs, which were previously known to undergo an Rrp6p-dependent trimming/processing of their 3’-extended precursor intermediate (32, 36, 87–90), displayed a little and insignificant amount of accumulation (only 2-3 fold vs. 20-130 fold for the mature species) in the *rrp6*-Δ strain. We believe that these proximal precursor amplicons exhibit minimal accumulation in the *rrp6-*Δ strain due to their Rrp6p-dependent trimming. All of these findings suggest the existence of a novel Rrp6p-dependent nuclear decay of the mature and polyadenylated version of small non-coding RNAs, e.g., 5S, 5.8S, snRNAs, and select snoRNAs. This conclusion was further correlated to the rate of the decay of these mature and precursor amplicons of these rRNA species in WT and *rrp6-*Δ strains (see below).

### Enhancement of the steady-state levels of the mature polyadenylated form of the small non-coding RNAs is directly associated with their diminished decay rates in *rrp6-*Δ strain

To further corroborate if the steady-state enhancement of the mature polyadenylated pool of 5S, 5.8S rRNAs, sn-, and select snoRNAs in *rrp6*-Δ strain is associated with their nuclear decay, we determined the decay rates of polyadenylated versions of some of these mature and 3’-extended precursor forms of various small ncRNA species in the wild-type and *rrp6*-Δ strain by shutting of the RNA polymerase I, II and III transcription. The decay rates of the mature form of 5S and 5.8S rRNA pools (Fig. 6A and B) displayed a significantly diminished decay rate in the *rrp6*-Δ strain with a concomitant increase of their half-life values by about 4-5 fold (from 107 minutes and 99 minutes in the wild-type to 190 minutes and 395 minutes in *rrp6*-Δ strain for total mature 5S and 5.8S rRNA respectively). Remarkably, the decay rates of 3’-extended pre-5S precursor species did not reveal any alteration of decay rate and half-life values in these strains (Table 1). Moreover, analysis of the decay rates and half-life values of the pre-5.8S rRNA precursor, however, indicated that the pre-5.8S rRNA-I (representing the 5.8S + 30 intermediate) displayed a ≈2.5 fold enhancement (from 108 minutes in WT to 265 minutes in *rrp6*-Δ strain), and pre-5.8S rRNA-II demonstrated a very modest stabilization in *rrp6*-Δ strain approximately 1.6-fold (from 180 minutes in WT to 298 minutes in *rrp6*-Δ strain) thereby corroborating that the difference in the steady-state level enhancement of the mature polyadenylated forms of these rRNA species is associated with their decay rates that are dependent on Rrp6p (Figs. 6A and B). Thus, the decay rates and half-life values of various mature and precursor species of these small rRNAs are extremely well-correlated in wild-type and *rrp6*-Δ strain. Although the measured values of steady-state and half-life of total 5.8S rRNA comprise a small contribution of the steady-state and half-life values of pre-5.8S rRNA-I species (the 5.8S+30 intermediate), our data strongly indicate that mature and polyadenylated forms of 5S and 5.8S undergo an Rrp6p-dependent nuclear decay.

**Table 1:** Half-life values of various non-coding RNAs in WT (*RRP6*+) and *rrp6*-Δ yeast strains (in minutes)

**Figure 1: Polyadenylated version of mature forms of low molecular weight ribosomal RNAs accumulate**
| ncRNAs | 5S | pre 5S | 5.8S | pre 5.8S I | pre 5.8S II | SNR10 | pre snR10 A | SNR13 | pre snR13 A | pre snR13 B |  |  |  |  |
| --- | --- | --- | --- | --- | --- | --- | --- | --- | --- | --- | --- | --- | --- | --- |
| WT | 107.4 | 374.14 | 98.9 | 108 | 179.93 | 29.38 | 376.57 | 33.19 | 342.66 | 362.12 |  |  |  |  |
| *rrp6Δ* | 190.6 | 358.28 | 395.05 | 265.23 | 298.51 | 355.62 | 454.93 | 479.50 | 531.1 | 564.04 |  |  |  |  |
| ncRNAs | U1 | Pre U1 A | Pre U1 B | U2 | Pre U2 A | Pre U2 B | U4 | Pre U4 A | Pre U4 B | U5 | Pre U5 A | Pre U5 B | U6 | Pre U6 A |
| WT | 14.71 | 225.1 | 360.5 | 18.05 | 239.48 | 307.17 | 85.28 | 214.71 | 323.27 | 25.17 | 273.91 | 267.82 | 101 | 340.61 |
| *rrp6Δ* | 414.64 | 356.9 | 372.9 | 236.1 | 315.05 | 429.06 | 352.66 | 334.48 | 428.02 | 232.75 | 241.99 | 253.39 | 479.58 | 447.7 |

**Figure 6:**
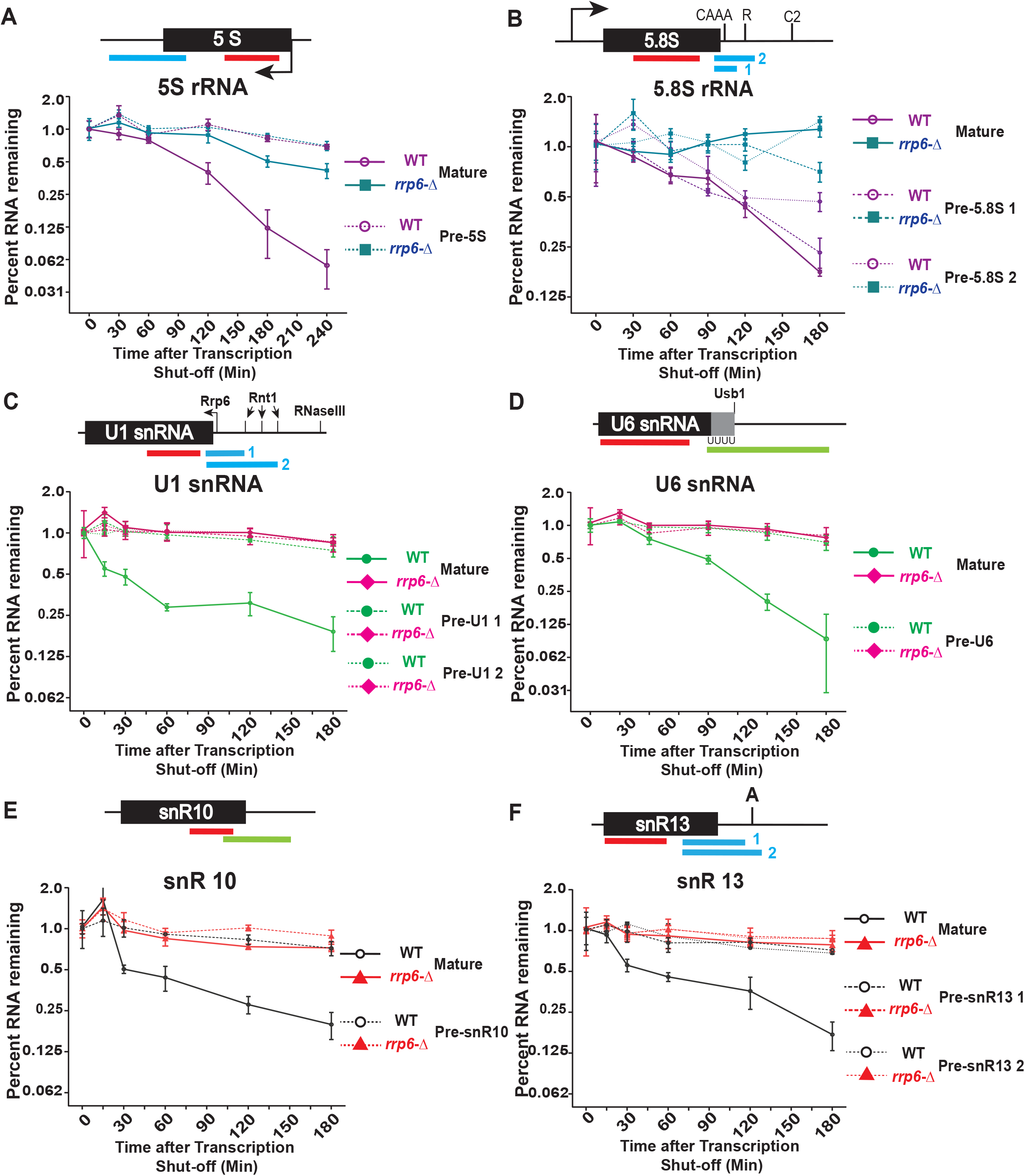
Transcription Shut-off experiment showing polyadenylated mature forms of the small non-coding RNAs undergo an active degradation by Rrp6p. Time kinetics of the steady-state levels of total and precursor small non-coding RNAs following the transcription shut-off revealing their decay rates in wild-type and *rrp6*-Δ strains. The decay rates were determined from four independent experiments (biological replicates, n=4) by qRT-PCR analysis (using amplicons indicated in thick red/blue/green lines below each gene sequence), and the signals were normalized to either *SCR1* RNA (in case of inhibition of RNA Pol-II) or *ACT1* mRNA (in case of inhibition of RNA Pol-I and III). Normalized signals (mean values ± SD) were presented as the fraction of remaining RNA (relative to normalized signals at 0 min) as a function of time of incubation in the presence of either transcription inhibitors BMH-21 (for RNA Pol I Transcripts); 1,10-phenanthroline (for RNA Pol II Transcripts); and ML-60218 (for RNA Pol III Transcripts). WT and *rrp6-*Δ are denoted with violet/blue, green/pink, and black/red colored lines for rRNA, snRNA, and snoRNA. qRT-PCR assays, Total RNA/cDNA isolations were carried out as described in materials and methods.

The decay rate (Fig. 6 and supplementary Fig. S2) and half-life of the mature and polyadenylated forms of all the snRNAs and two select snoRNA species, snR10 and snR13 (Table 1 and 2) unequivocally indicated that their decay becomes substantially diminished in the *rrp6*-Δ strains with a concomitant increase in their half-life values (12 to 14 fold enhancement for mature polyadenylated forms of snoRNAs, snR10 and snR13 and 5-28 fold increase for various snRNAs) (Fig. 6 and supplementary figure S2, Table 1). Interestingly, a comparison of the decay rates and half-life values of all the polyadenylated 3’-extended precursor forms of all the snRNAs and two snoRNAs very clearly revealed that they all displayed a minor and statistically insignificant amount enhancement (1-3 to 1.5 fold increment for different sn- and snoRNAs). Collective data is thus consistent with the conclusion that mature 5S, 5.8S rRNAs, sn- and select snoRNAs are subject to an active nuclear degradation by Rrp6p.

### Nuclear decay of mature small non-coding RNAs requires Rrp47p and both canonical and non-canonical poly(A) polymerase Pap1p- and Trf4p-dependent polyadenylation

The nuclear form of the exosome specifically remains associated with Rrp6p (15, 42) and several ancillary factors, Lrp1p/Rrp47p (specific to nuclear exosome, CID in humans), Mpp6p (specific to nuclear exosome) (43, 86, 91, 92). Interestingly, Rrp6p alone is also known to interact with Rrp47p *in vitro* (93) and *in vivo* (76, 94) independent of the core nuclear exosome. Moreover, depletion of the Rrp47p resulted in the accumulation of Rrp6p-specific processing and degradation intermediates/precursors (94). Moreover, Rrp6p and Rrp47p were reported to stabilize each other (91, 95), and deletion of either Rrp6p or Rrp47p in the strains additionally lacking Mpp6p leads to the synthetic lethality, thus indicating that these factors functionally interact with one another (92). Thus, the physical association of Rrp47p and Mpp6p with Rrp6p and the previous instance of the assistance of Rrp47p/Mpp6p in the function of Rrp6p prompted us to evaluate if Rrp6p-dependent degradation of the small non-coding RNAs also requires Rrp47p and Mpp6p.(94)

Having shown that Rrp6p targets the mature polyadenylated forms of the small non-coding RNAs, we address which ancillary factors are vital for this activity. Towards this, we first addressed if the ancillary cofactors of Rrp6p, Rrp47p, and Mpp6p are involved in this nuclear decay. Consequently, we determined the levels of all of these mature RNA species in yeast strain carrying a deletion in *RRP47* and *MPP6* and deletion in both of these genes using both random-primed and oligo-dT-primed cDNA samples from these strains. As shown in figure 7A-H, all of these ncRNAs, 5S,5.8S, U-snRNAs (data for U5 not shown) and snoRNAs, snR10, and 13 displayed a dramatic enhancement of their steady-state levels in *rrp6-*Δ, *rrp47*-Δ, and *rrp6-*Δ *rrp47*-Δ double mutant yeast strains (Fig. 7A-H). No such enhancement in the levels of any of these mature RNA species was noted in yeast strain carrying *mpp6*-Δ deletion (Fig. 7A-H), suggesting that Mpp6p is not involved in this nuclear decay. The exonuclease domain of Rrp6p is vital for this nuclear decay is demonstrated by a dramatic and comparable enhancement (≈20-150) of the mature and polyadenylated version of these ncRNAs in a yeast strain with *rrp6-3* allele harboring a catalytically inactive point mutation (D238A) in the exonuclease I domain of Rrp6p (96), thus unequivocally affirms that the destruction of its exonucleolytic activity accompanies their steady-state level enhancement in *rrp6*-Δ strain. Remarkably, the extent of enhancement of the steady-state levels of all these ncRNAs in double mutant *rrp6-*Δ *rrp47*-Δ strain is no better than their levels found in each of the *rrp6-*Δ, and *rrp47*-Δ single mutant strains, thereby exhibiting genetic epistasis between *RRP6* and *RRP47* genes concerning the nuclear decay of these ncRNAs. This data strongly supports the idea that both Rrp6p and Rrp47p are acting together in this nuclear decay pathway to target these mature and polyadenylated ncRNAs. To further corroborate the finding that this Rrp6/47p-dependent decay of ncRNA is independent of the nuclear exosome activity, we also show Rrp6p and Rrp47p interact with each other *in vivo* by Co-IP using Anti-TAP antibody from a yeast strain that expresses Rrp47-TAP (Fig. 7I). Moreover, we also showed Rrp6p co-purifies with Rrp47p as a separate complex from the exosome (Exo-11) in a gel exclusion column chromatographic procedure (Fig. 7J), thereby indicating that this independent exonuclease complex catalyzes this nuclear decay. Thus, our data strongly suggest that Rrp6p and Rrp47p constitute a separate and exosome-independent nuclear RNA degradation system that targets the polyadenylated version of mature small non-coding RNAs.

**Figure 7:**
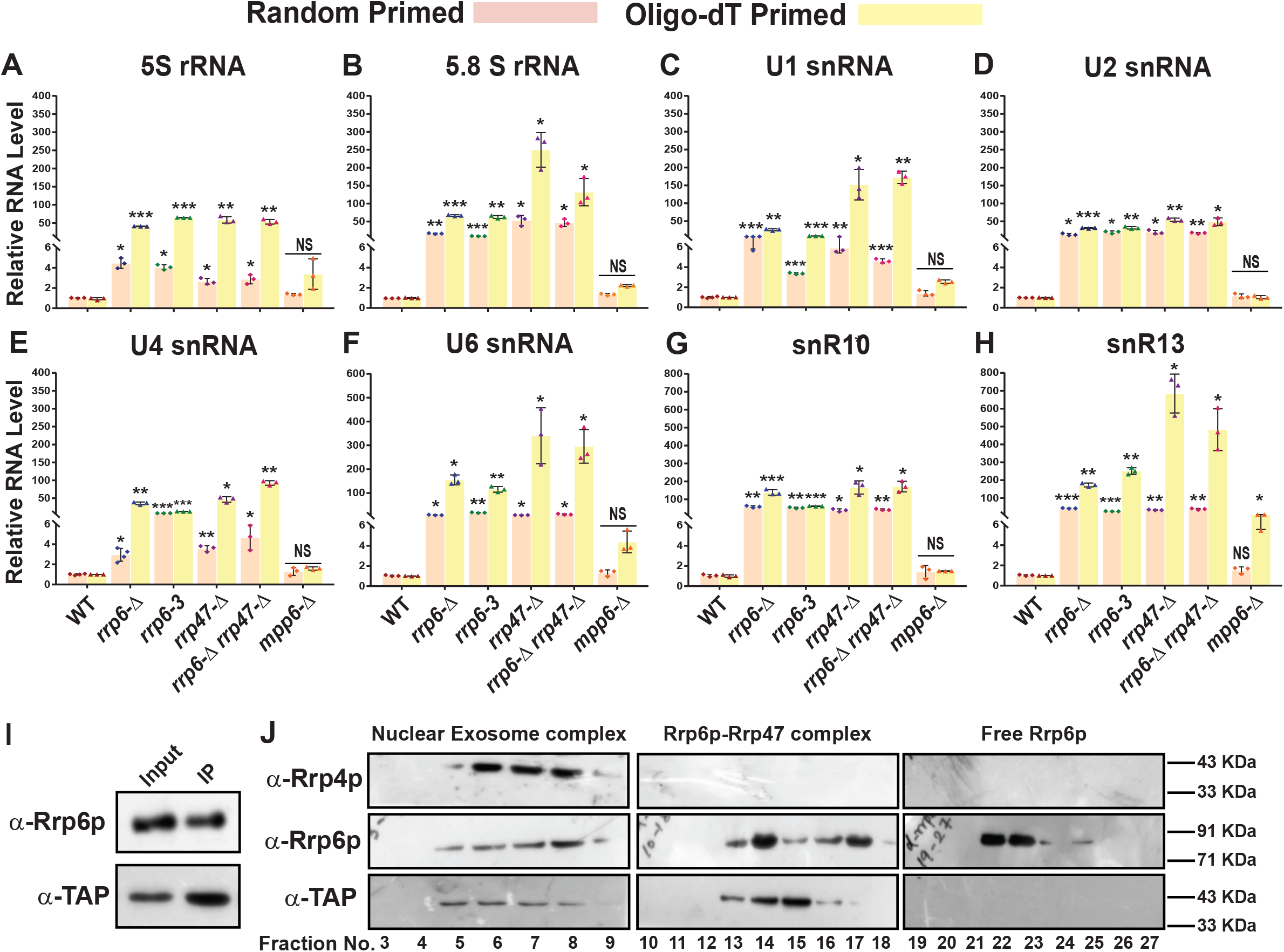
Rrp6p-dependent degradation of the mature small non-coding RNAs requires its functional exonuclease domain and ancillary nuclear factor Rrp47p/Lrp1p, but does not require Mpp6p. (A-H) Scattered/Bar plot revealing the steady-state levels of mature forms of various small ncRNAs estimated by qRT-PCR from the 2 ng cDNA samples prepared using either random hexanucleotide primers (orange bars) or oligo-dT_30_ anchor primer (yellow bars) from the indicated yeast strains carrying mutations in the *RRP6* (*rrp6-*Δ*, rrp6-3*)*, RRP47* (*rrp47*-Δ), *MPP6* (*mpp6-*Δ) genes and *rrp6-*Δ *rrp47*-Δ double mutant alleles together*. SCR1* (in the case of Random Primer) and *ACT1* mRNA (in the case of Oligo dT Primer) were used as the internal loading control. Normalized values of each of the ncRNAs in the wild-type yeast strain were set to 1. Three to Four independent cDNA preparations (biological replicates, n = 3, in some cases 4) were used to determine the levels of various ncRNAs. The statistical significance of difference reflected in the ranges of P values estimated from Student’s two-tailed t-tests for a given pair of test strains for every message are presented with the following symbols, *<0.05, **<0.005, and ***<0.001; NS, not significant. qRT-PCR assays, Total RNA/cDNA isolations were carried out as described in materials and methods. (I-J) Co-IP and fractionation/separation profile of the nuclear exosome and Rrp6/47p in Biogel P-200 gel-filtration chromatographic column showing Rrp6p and Rrp47p acts as a complex independent of core nuclear exosome. For (I), total cellular protein extract was prepared from the yeast strain expressing Rrp47-TAP followed by co-immunoprecipitation using TAP antibody. Immunoprecipitate was further tested with the anti-Rrp6p antibody for the presence of Rrp6p. For (J), the same Rrp47-TAP protein extract was fractionated by Biogel P-200 Gel filtration column, and the eluate fractions were subjected to western blotting analysis using anti-Rrp4p, anti-Rrp6p, and anti-TAP (For detection of Rrp47p). The Protein isolation, western blotting, and Co-IP were carried out following standard protocols described in materials and methods.

Further, we addressed which cellular polyadenylation activity is required to make the small non-coding RNAs’ polyadenylation marks. Two different poly (A) polymerases are present in the genome of the baker’s yeast *Saccharomyces cerevisiae*. The major canonical poly(A) polymerase is Pap1p (97), an integral part of the cleavage/polyadenylation complex 98, which catalyzes the independent template addition of poly(A) tails most messenger RNAs. We first address if Pap1p is playing any role in the polyadenylation of these mature form of small ncRNAs. Consequently, we determined if any alteration of the steady-state levels of the mature form of the small ncRNAs in the yeast strain carrying a temperature-sensitive (ts) *pap1-1* allele (98) and compared their steady-state levels estimated from cDNAs primed with both random and oligo-dT primers in wild-type and yeast strains carrying, *rrp6-*Δ, *pap1-1* and *pap1-1 rrp6-*Δ double alleles at the permissive and non-permissive temperature of 25°C and 37°C. As shown in Figure 8, the steady-state levels (both the total as well as polyadenylated fractions) of none of these small ncRNAs displayed any enhancement in *pap1-1* strains at the permissive condition of 25°C (Fig. 8A-F, left histogram) as the mutant phenotype is not manifested in this condition. However, at non-permissive condition of 37°C an enhancement of the levels of these ncRNAs in the *pap1-1* and *pap1-1 rrp6*-Δ double mutant strains determined only from random-primed cDNA were observed, which is similar and comparable to that observed in single *rrp6-*Δ strains (Fig. 8A-F, right histogram). Strikingly, the steady-state levels of all of the ncRNAs were found to be dramatically declined in strains carrying *pap1-1* and *pap1-1 rrp6*-Δ alleles, which were determined from oligo-dT-primed cDNA (Fig. 8A-F, right histogram). Although, initially this finding appeared unexpected, this findings makes perfect sense since at 37°C in *pap1-1* strain, the global RNAs become unadenylated owing to the complete manifestation of the *pap1-1* phenotype (98) and thus oligo-dT primers failed to prime gross cDNA synthesis under this condition leading to very low qRT-PCR signal. Thus, our results thus strongly favor the argument that Pap1p is catalyze the polyadenylation of these small non-coding RNAs, making the mark of degradation. Notably, aberrant/faulty mRNA and a few non-coding RNA substrates in the nucleus and nucleolus were also known to undergo short polyadenylation by the TRAMP4/5 complex before targeted for decay by the nuclear exosome (30, 66, 99). As shown in Fig. 1 and 2, our initial results suggest that the steady-state levels of the small non-coding 5S and 5.8S rRNAs, snRNAs, and select snoRNAs were very marginally augmented in yeast strain carrying the *trf4-*Δ alleles (Figs. 1 and 2). A more careful analysis of both the random-primed and Oligo-dT-primed cDNA samples prepared from isogenic *trf4-*Δ and *trf5-*Δ strains displayed a moderate accumulation of only snRNAs and select snoRNAs in *trf4-*Δ strain, which is comparable to their accumulation in *rrp6-*Δ strain (Fig. 8I-N). Interestingly, no significant increment of 5S and 5.8S rRNA were noted either in *trf4-*Δ or in *trf5-*Δ strain (Fig. 8G-H). Thus, these data strongly suggest that Pap1p carries out polyadenylation of the mature small rRNA species and that of other small ncRNAs is carried out by both Pap1p and Trf4p, in which significant contribution comes from Pap1p, and Trf4p plays a minor functional role (see discussion).

**Figure 8:**
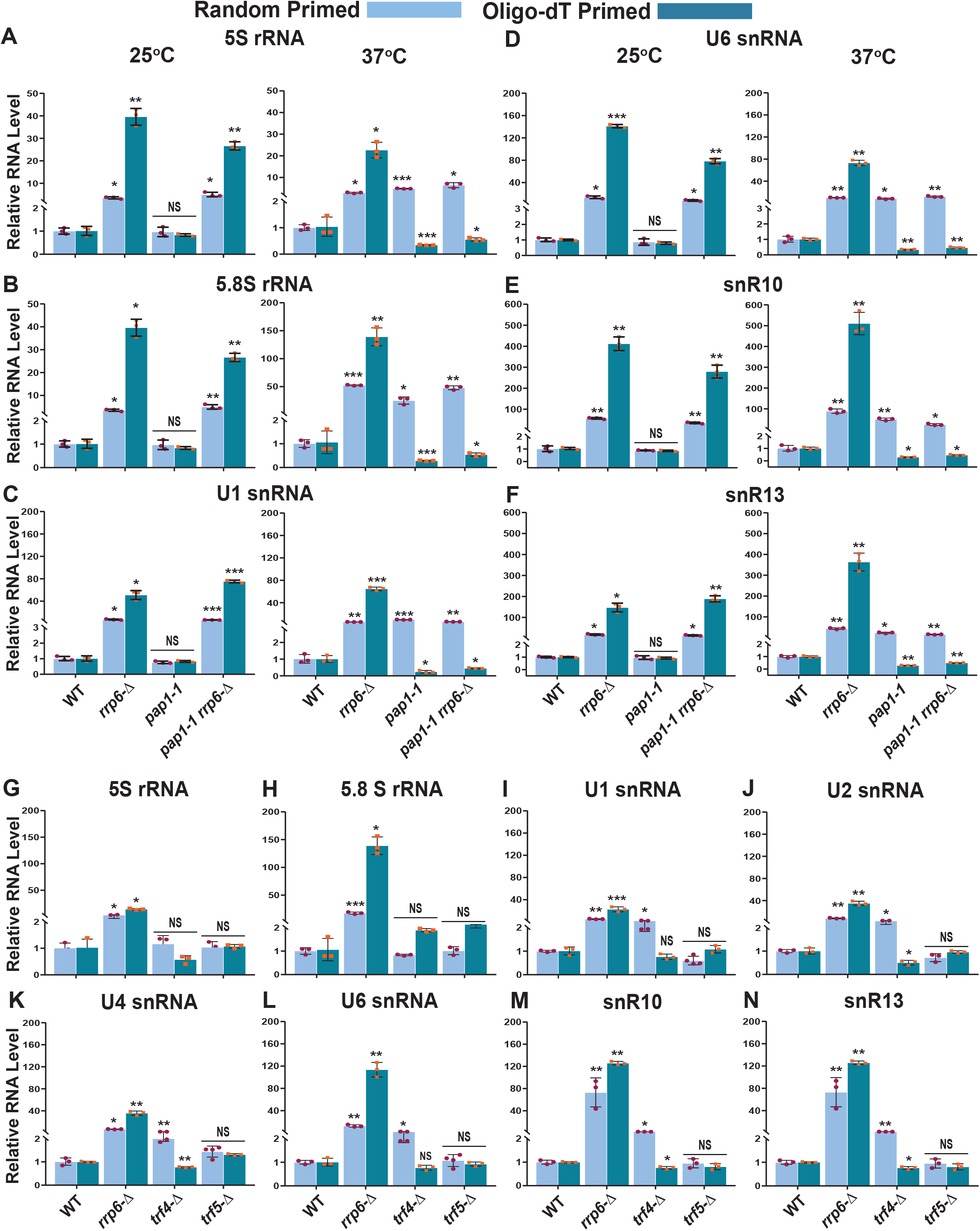
Both the canonical Poly(A) polymerase Pap1p and non-canonical Poly(A) polymerase Trf4p play a vital role in the polyadenylation of the smaller ncRNAs. Scattered/Bar plot revealing the steady-state levels of various low molecular weight ncRNAs estimated from the 2 ng cDNA samples prepared using random hexanucleotide primers (sky blue bars) or oligo-dT_30_ anchor primer (indigo blue bars) by qRT-PCR from WT and yeast strains carrying mutations in the *RRP6* (*rrp6-*Δ)*, PAP1* (*pap1-1*), *pap1-1 rrp6-*Δ double mutant alleles (panels A-F) or WT, *RRP6* (*rrp6*-Δ), *TRF4* (*trf4-*Δ), *TRF5* (*trf5-*Δ) genes (panels G-N). The *pap1-1* and *pap1-1 rrp6-*Δ strains were pre-grown at 25ºC, followed by splitting the culture into two halves. One half was continued to grow at 25°C for 12 hours, and a 12h shift to 37ºC were performed to the other half of the culture before harvesting them. Total RNA, cDNA isolation from them followed by qRT-PCR reaction carried out as described in materials and methods. *SCR1* (in the case of Random Primer) and *ACT1* mRNA (in the case of Oligo dT Primer) were used as the internal loading control. Normalized values of each of the ncRNAs in the wild-type yeast strain were set to 1. Three independent cDNA preparations (biological replicates, n = 3) were used to determine the levels of various ncRNAs. The statistical significance of difference reflected in the ranges of P values estimated from Student’s two-tailed t-tests for a given pair of test strains for every message are presented with the following symbols, *<0.05, **<0.005, and ***<0.001; NS, not significant.

### Genome-wide analysis of non-coding RNAs shows a dramatic accumulation of transcripts corresponding to the mature region of the select snoRNA genes in *rrp6-*Δ strain

To further corroborate our finding of differential and dramatic accumulation of mature species of snoRNAs presented above, we employed a previously published database (accession number GSE135056) (100) describing the annotations of yeast transcripts and performed a differential expression analysis in the genome-wide scale using normalized RNA-sequencing reads by EdgeR (101–103) in the WT and *rrp6-*Δ strains. The annotations used for our study included 82 snoRNAs. Interestingly, in *rrp6-*Δ strain, the majority of the 82 snoRNA genes were found to undergo upregulation by more than two-fold (LogFC change cutoffs = +/− 1, p-value cutoff = 0.05, Fig. 9A-C). To further evaluate which part of the snoRNA gene accumulates the maximum sequence reads, we plotted the number of reads aligned per base pair for four different snoRNAs, snR10, snR13, snR48, and snR71 across their genomic loci as well as along their 3’-extended regions in both WT and *rrp6-*Δ yeast strain (Fig. 9D-G). To critically analyze these data, we plotted the read-accumulation in their genomic loci at two different zoom-levels, one distant view to accommodate the 3’-extended regions (top panels) and a close-up view (bottom panels). A careful comparison of the sequence read-data for these snoRNAs reveals that all of them have increased accumulation of reads in the *rrp6-*Δ strain, and the majority of the read accumulation occurs in the mature region of these snoRNA genes. Although a substantial aggregation of reads was found in the extended region of some of them, many of these genomic loci also harbor either another mRNA ORF, which is overlapping with the snoRNA gene (such as snR10 overlapping with YGL088W) or has another mRNA ORF in the vicinity with opposite orientation (such as snR71 has *LIN1* in its 3’-side with opposite orientation concerning the direction of transcription). The presence of these mRNA genes in the overlap/adjacent regions of snoRNA genes might contribute to the observed accumulation of sequence reads at the 3’-extended regions of some of these snoRNAs, owing to the possible presence of potential antisense transcripts (strong targets of Rrp6p). Although this made the analysis and interpretation of the data a little complicated, even with a cautionary note in mind, we favor the view that the numerical value of the accumulated reads in the mature region of each of these snoRNA genes is substantially higher than that detected in their mature regions in the WT strain. Our analysis of the existing datasets thus clearly demonstrates that these snoRNAs’ mature RNA species are targeted by Rrp6p, thereby corroborating our finding from the experiments presented above.

**Figure 9:**
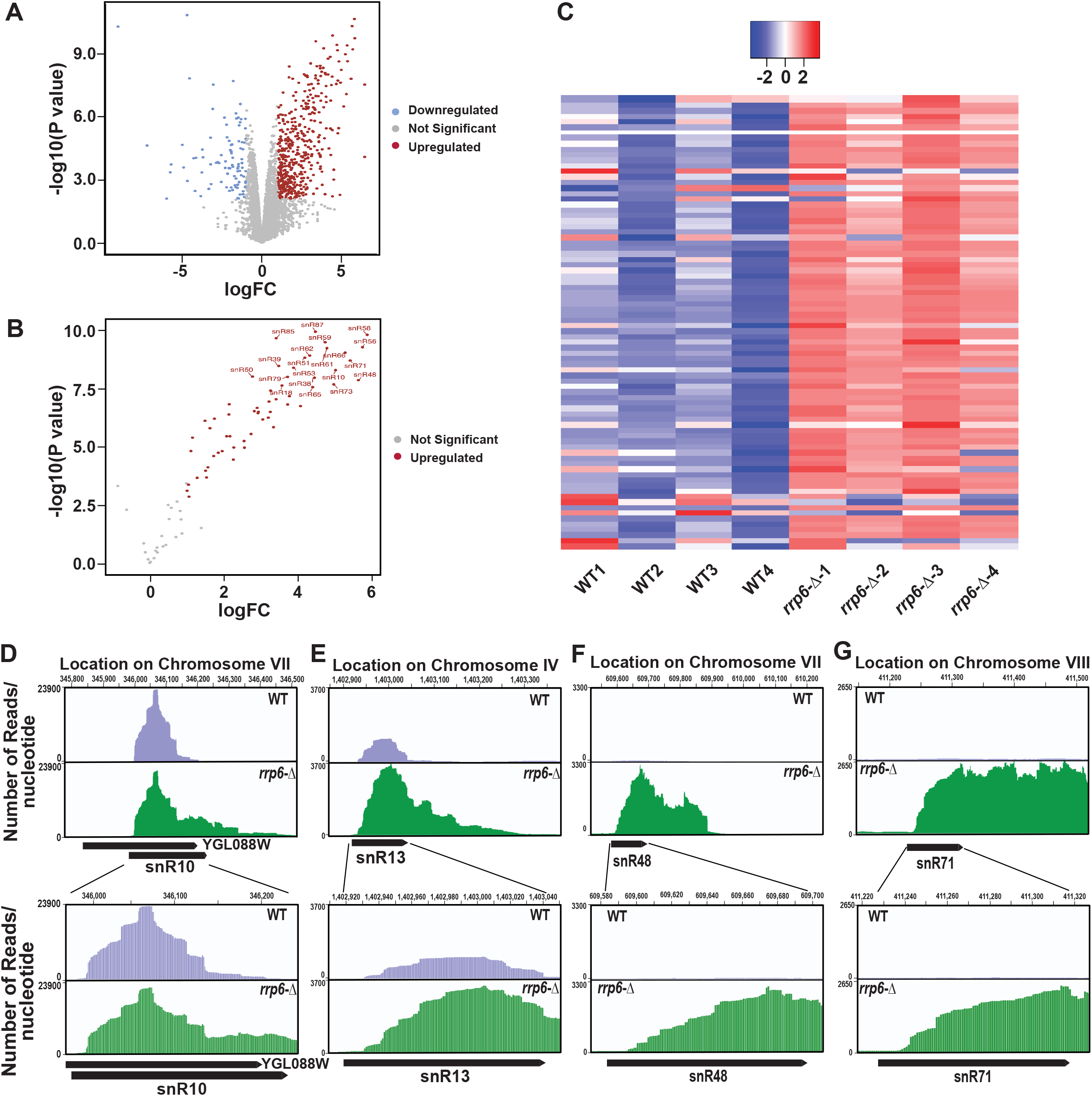
Analysis of a previously submitted RNA-seq data (Accession Number GSE135056) revealed a dramatic accumulation of reads corresponding to the mature region of several small nucleolar RNAs. Volcano plots depicting the differential expression of all annotated sequences (A) and all 82 sn- and snoRNA transcripts (B). (C) Heat Map of normalized counts, showing the expression pattern for all 82 sn- and snoRNAs across four independent biological replicates of WT and *rrp6-*Δ yeast strains. (D) Graphical representation showing the relative amount of reads mapped to the genomic locus corresponding to small nucleolar RNAs snR10-YGL088W, snR13, snR48, and snR71. The top panels depict the distant view (accommodating the 3’ extended regions), and the bottom panels show the close-up views for each snoRNA loci. The location of transcripts and the direction of transcription are shown below the graph (drawn in scale) by the solid black arrow-headed rectangles.

## Discussion

The processing events of various non-coding RNAs involve both the endonucleolytic cleavage and processive exonucleolytic trimming from 5’- and 3’-termini (1–7)^,8–19^. The nuclear exosome, consisting of nine catalytically inactive and two active subunits, plays a direct functional role in their 3’-end processing and indirectly impacts their covalent modification events (15, 41, 52, 104, 105). Although traditionally all the exosome subunits were believed to collectively participate in all the exosomal functions 8,11,41,52–64, a body of evidence indicates that many subunits of the exosome may physically exist and function as a separate entity independent of the ‘core’ exosome (30, 44, 72, 64–71). In this work, we unveiled a novel functional role of vital exosomal component, Rrp6p and its cofactor Rrp47p in the degradation of mature polyadenylated versions of several small non-coding RNAs, including 5S, 5.8S rRNAs, all sn- and many snoRNAs in the nucleus of baker’s yeast *Saccharomyces cerevisiae*. Our data showed that mature form of these ncRNAs undergo rapid degradation in Rrp6p-Rrp47p dependent manner and requires prior polyadenylation catalyzed either by Pap1p (in case of 5S and 5.8S rRNAs) or jointly by Pap1p and Trf4p (in case of sn- and snoRNAs). Notably, Rrp6p is involved in the mature 3’-end formation of many snoRNAs and is believed to trim the last few nucleotides to form mature 3’-end of these snoRNAs (85). Therefore, our finding of the Rrp6p-dependent degradation of mature species of 5.8S rRNA, snRNAs, and several snoRNAs may be viewed and interpreted as a consequence of the 3’-end trimming reaction of the 5.8S+30 intermediate, and the 3’-extended precursor species of several sno- and snRNAs by Rrp6p (41, 52, 85, 88, 105–107) and not as the degradation of their mature forms. However, a careful analysis as carried out by precise measurements of various mature and precursor fragments using specific amplicons described in the results section above enabled us to conclude with a high degree of certainty that Rrp6p along with Rrp47p targets and degrades the mature polyadenylated forms of these small ncRNAs besides trimming their 3’-extended precursors in some cases as reported before (41, 52, 85, 88, 105–107). This conclusion was also obvious from previous data published from many laboratories. In many of those studies addressing the abundance of 5.8S and its 3’-extended precursor 5.8S+30 intermediate using northern blot analysis, it was observed that in wild type (*RRP6*^*+*^) strain only the mature 5.8SS/L is visible and in *rrp6-*Δ strain two bands are detectable, one corresponding to the 5.8SS/L band (whose intensity is either equal to or slightly less than the corresponding intensity in WT cells) and the other corresponding to 5.8S+30S/L intermediate (79, 85, 94, 96, 108, 109). From these data Rrp6p was concluded to participate only in the exonucleolytic processing/trimming of the last 30 nucleotides of 5.8S rRNA. In most of these studies, no potential role of Rrp6p to target and degrade the mature 5.8SS/L rRNA was concluded. We would like to reconcile these earlier observations and our present data by arguing that Rrp6p targets and degrades the mature polyadenylated version of 5.8S rRNA besides acting on the 5.8S+30 intermediate rRNA. The presence of the mature 5.8SS/L band in the *rrp6-*Δ strain in equal or slightly less intensity in addition to newly appeared the 5.8S+30S/L band in those earlier observations from the northern analysis completely support our observation and justifies our conclusion stated above. We would like to argue that if Rrp6p would target and trim only the 5.8S+30S/L precursor intermediate and not the mature 5.8SS/L RNA, then in the northern analysis the band corresponding to the mature 5.8SS/L would disappear and would shift to the 5.8S+30S/L position, as it happens for the some of the mature and 3’-extended precursor species of U24, snR40 and snR72 RNAs (96), U14, U18, and snR38 (94). Interestingly, some of these studies also revealed that the relative abundance profiles of several non-coding RNAs, snR13 (94, 110, 111), snR33 (110) U4 snRNA (105) snR8, snR14, snR77, snR44 snoRNAs (94) and their 3’-extended precursor species in WT and *rrp6*-Δ strain displayed a similar relative abundance profile of 5.8SS/L and 5.8S+30S/L intermediate, thereby suggest that Rrp6p and Rrp47p targets and degrades their mature species in addition to their 3’-extended precursor. Thus, a critical comparative analysis of our present data and the observations reported in the earlier studies leads to the conclusion that Rrp6p/47p targets and degrades the polyadenylated mature species of 5S, 5.8S, all snRNAs and several select snoRNAs. Consistent with this conclusion, analysis of the decay rates of the mature and one or more 3’-extended precursor of each of the 5S, 5.8S, snRNAs and selected snoRNAs (Fig. 6) and subsequent comparison of their half-life values (Table 1) revealed that the mature forms each of these non-coding RNAs are undergoing rapid decay in WT yeast strain in Rrp6p-dependent manner. Loss of Rrp6p in the same isogenic strain resulted in their diminished decay rates with concomitant and dramatic enhancement of their half-life (Table 1). Subsequent analysis of previously published genome-wide RNA-sequencing datasets (100) and the resulting differential expression profile of the snoRNAs in WT and *rrp6-*Δ strains revealed that almost all the test 82 snoRNAs were significantly up-regulated in *rrp6-*Δ strain (Fig. 9A-C). Further analysis showed that accumulation of normalized sequence reads were principally detected in the mature region across the genomic loci of four test snoRNAs, snR10, snR13, snR48 and snR71 (Fig. 9D-G), thus supporting the conclusion that the mature forms of these small non-coding RNAs were targeted by Rrp6p/47p.

Remarkably, our finding of accumulation of mature and polyadenylated forms of small ncRNAs is reminiscent of a previous study, demonstrating that a discrete domain rich in polyadenylated RNA appears in the yeast strain lacking functional Rrp6p (112). This study also showed that the polyadenylated transcripts accumulate in a distinct site in the nucleolus called poly(A) domain, which is different from the sites of many of these ncRNAs’ maturation and also require Rrp6p, Pap1p, and Trf4p for this poly(A) specific foci formation. Other than Rrp6p, Pap1p, and Trf4p, however, depletion of the nuclear exosome components, Rrp41p, and TRAMP component Mtr4p, also led to the formation of sub-nucleolar poly(A) domain formation (112). However, in our experimental system contribution of any core exosomal component (except Rrp6p) and TRAMP component (except Trf4p) was not detected. Interestingly, although the nucleolar poly(A) domain was found to be enriched with polyadenylated and 3’-extended precursor form of U14 snoRNA and possibly their mature form, this study did not evaluate what other kinds of RNAs are present in such poly(A) domain. Neither it was tested if the loss of Rrp47p leads to the poly(A) domain formation in the nucleolus (112). Nevertheless, it remains to determine the relationship between the poly(A) domain as reported earlier (112) and the dramatic accumulation of polyadenylated forms of mature small non-coding RNAs presented here.

Our demonstration of co-purification of Rrp6p and Rrp47p as an exosome-independent and separate protein complex is in good agreement with our observation of the requirement of only these two factors for the degradation of the mature polyadenylated forms of the small non-coding RNAs. However, it remains to evaluate if the Pap1p and Trf4p is also part of the same complex or exert their effect independently of this complex. Although the complete protein and RNA composition of the Rrp6-Rrp47p complex are yet to be revealed, it is reasonable to state that this complex may be responsible for carrying out the degradation of polyadenylated mature non-coding RNAs in the nucleolus and also other exosome-independent functions reported earlier. Future work will throw more light on this unresolved issue.

Although both the known poly(A) polymerases, Trf4p and Pap1p, were found to play an essential role in polyadenylating the small non-coding RNAs, their fundamental feature of catalysis is quite different. Pap1p is principally responsible for adding long poly(a) tails to the nuclear pre-mRNA precursors in a processive manner, and the speed of the reaction is breakneck and efficient. In contrast, the rate of catalysis by Trf4p is very slow and inefficient, and it adds only a few residues of adenine in a distributive manner (30). The evidence of the involvement of Pap1p in the polyadenylation of these small non-coding RNAs strongly suggests that the average length of these small ncRNAs’ tails is reasonably long, which is also supported by the fact that these RNAs could be efficiently primed by oligo-dT12-18 primer. Although the sequence of their participation in polyadenylating these RNAs is not verified experimentally, it is reasonable to suggest that Trf4p participates first to form a short oligo tail of 4-6 nucleotides used as a primer for the subsequent polyadenylation by Pap1p. Consistent with this assumption decreased polyadenylation of these small non-coding RNA was observed in a double *pap1-1 rrp6-*Δ mutant strain compared to the *rrp6*-Δ mutant (Fig. 8, A-F).

Although this observation’s physiological significance is uncertain at this point, this phenomenon could be related to regulating the speed and efficiency of ribosome biogenesis during the stress response. As ribosome-biogenesis is an incredibly energy-requiring process, a cell would likely prefer to speed up the ribosome biogenesis process during a favorable environment and growth. During any stress, the biogenesis of ribosomes would likely be restricted and regulated. Although the exact molecular mechanism is not known, it might not be unreasonable to invoke that perhaps during stress, this exosome independent Rrp6p/47p dependent degradation of small non-coding RNAs could serve as a major mechanism to regulate and limit ribosome biosynthesis, which in turn could induce the proteolysis of the ribosomal proteins (113). Recent demonstration of rRNA processing inhibition during Integrated Stress Response (ISR) in mammalian cell (114) supports this hypothesis. However, further experiments addressing and requiring to resolve this issue and confirm this hypothesis is currently underway.

## Methods

### Nomenclature, yeast strains, media, and yeast genetics

Standard genetic nomenclature is used to designate wild-type alleles, as *CBC1*, *RRP6*, *RRP4*, *TRF5,* etc. Mutant alleles that produce defective gene products and hamper the functions of the particular gene are designated *rrp4*, *rrp6, dob1 or pap1* followed by the allele number, e.g*., rrp4-1,rrp6-3, dob1-1, pap1-1* whereas *rrp6-∆, cbc1-∆ or rrp47::KANMX, his3::hisG, tif4631::URA3* gene symbols designate deletion or disruption in *RRP6, CBC1, RRP47, HIS3,* and *TIF4631* genes respectively. The protein encoded by a specific gene is denoted, for example, Cbc1p, which is encoded by *CBC1*. Standard media like YPD, YPGal, YPRS, and other omission media like URA dropout, TRP-dropout, and so on were used to propagate and maintain yeast. Yeast genetic analysis was carried out by standard procedures as described before (115). All the yeast strains are listed in Table S1 (Supporting Information).

### Plasmids and oligonucleotides

The plasmids were either procured from elsewhere or were constructed in this laboratory as described previously (116). Cloning experiments were performed for creating different disruptor plasmids for genes like *TRF4, TRF5, AIR1,* and *HIS3*. All the oligonucleotides were obtained commercially from Integrated DNA Technology (Coralville, IN, USA). The plasmids and oligonucleotides used in this study are listed in Tables S2, S3, and S4 (Supporting information), respectively.

### Construction of various deletion constructs in an isogenic series

An isogenic series of different strains deficient in CTEXT, the nuclear exosome complex, and TRAMP components were constructed by following standard recombinant DNA procedures.*cbc1-*Δ*, tif4631-*Δ*, upf1-*Δ*, rrp6-*Δ*, trf4-*Δ*, trf5-*Δ*, air1-*Δ*, rrp47-*Δ*, rrp6-*Δ *rrp47-*Δ, and *mpp6-*Δ strains were constructed using the one-step gene replacement method.

### Galactose induction of *GAL10::RRP41* gene

The native promoter of the *RRP41* gene in wild type strain is replaced by p*GAL1*/*GAL10* promoter by homologous recombination and confirmed using genomic PCR to get exosome deficient strain *HIS3*-*GAL10:: protA-RRP41*. This strain, harboring p*GAL1*/*GAL10::RRP41* was grown in non-inducible, non-repressible 2% raffinose sucrose medium (YPRS) till O.D_600_ = 0.6. Galactose was then added to the culture at 2% final concentration, and cells were grown till the late log phase, followed by harvesting of cells after changing the media to Glucose for 2 hours. Total RNA was isolated from the cells harvested as described above for further downstream analysis.

### Construction of WT and *rrp4-1* yeast strains by two-step gene replacement

Two-step gene replacement was carried out as described before (115). The WT (yBD 5) and *rrp4-1* (yBD 16) strains were transformed using standard Lithium Acetate Method with *HIS3* disruptor fragment of pBD 216 containing *his3:: hisG-URA3-hisG* and then screened for positive candidates. Then colonies were screened, and *URA3*-*hisG* was popped out in the next step by growing the yeast strain in standard 5-Fluoro-orotic Acid media (5-FOA) to get yBD263 and yBD285. The integration of the deleted allele was finally confirmed by genomic PCR using appropriate primer sets.

### RNA analyses and determination of steady-state of RNAs

As described earlier, the total RNA was isolated by harvesting respective yeast strains after growing them till the late log phase (OD_600_ 0.8-1). This is followed by the extraction of cell suspension in the presence of alkaline glass bead and phenol-chloroform–IAA (25:24:1). The RNA was finally recovered by precipitating it with RNAase-free absolute ethanol.

For cDNA preparation, 2μg RNA samples were treated with RNase free DNase I (Fermentas Inc.) at 37 °C for 30 min, followed by incubating it with 10 mM EDTA at 65 °C for 10 min. This is followed by cDNA synthesis by reverse transcription of 1μg of purified RNA by MMLV reverse transcriptase (BioBharati) for 50 min at 42°Cusing either random hexamers or oligo-dT_(12-18)_ primer.

### Real-Time qRT-PCR

Following the synthesis from total RNA samples, each cDNA was first quantified using Qubit® ds DNA-HS Assay Kit (Life Technologies, USA) following their recommendation. 2 ng of quantified cDNA from 3 to 5 biological replicate was used to quantify the levels of specific ncRNAs such as *rRNAs, snRNAs and snoRNAs snR10 and snR13*, by Real-time qPCR assays by using target-specific primers for mature and precursor regions and standard SYBR Green Technology using Power SYBR® Green PCR Master Mix (Applied Biosystems). qPCR assays were run in triplicate and conducted using an Applied Biosystems StepOneTM real-time PCR system to determine the absolute quantification of individual target mRNAs. For each target, ΔΔ-Ct method was used, and the abundance was determined from fluorescence by the StepOneTM software version 2.2.2 (Applied Biosystems).

### Analysis of stability of specific ncRNAs using Transcription Shut-off experiment

The stabilities of the various RNAs were determined by the inhibition of transcription with transcription inhibitors BMH-21 (for RNA-pol I Transcripts: rRNA 5.8s and ITS1), 1,10phenanthroline (for RNA-pol II Transcripts: mRNAs; snRNAs; and snoRNAs SNR10 and SNR13), and ML-60218 (for RNA-pol III Transcripts: rRNA 5s and snRNA U6) at required temperatures as described by (Das et al. 2000; Das et al. 2003;). Briefly, specific yeast strains were grown at 30°C till mid-logarithmic phase (OD_600_ 0.6) when transcription inhibitors BMH-21; 1, 10-phenanthroline; and ML-60218 were added to the growing culture at (25μM, 100 μg/ml, and 16μM) final concentrations respectively, followed by withdrawal of a 10 ml aliquot of growing culture at various time intervals after transcription shut-off, harvesting the cells and snap freezing the pellets in dry ice. It is followed by RNA isolation and determination of signals of various ncRNAs. The ncRNA levels were quantified from cDNA by real-time PCR analysis, and the signals associated with the specific ncRNA were normalized against *SCR1* signals. The decay rates and half-lives of specific mRNAs were estimated with the regression analysis program (Graphpad Prism version 7.04) using a single exponential decay formula (assuming mRNA decay follows first-order kinetics), y = 100e^−bx^ was used.

### Co-Immunoprecipitation

Cells were grown in 50ml YPD until OD_600_ 2.7-3.0 is reached. The cell lysate was prepared as described previously in 1ml of Buffer A. Protein A-^PLUS^ Agarose Beads (BioBharati LifeSciences Pvt. Ltd.) ~50ul bed volume per reaction were equilibrated twice with ten volumes of Buffer A. The beads for each washing was resuspended in Buffer A and kept on a rotator wheel for 5mins at 4°C. The beads were centrifuged at 3500rpm for 3 minutes each time to remove residual Buffer A. The extract was then pre-cleared by adding the equilibrated beads and incubation on a rotator wheel at 4°C for 30 minutes. The extract was then centrifuged at 3500rpm for 5minutes at 4°C to get rid of the beads. To the pre-cleared protein extract, 2.5ug of anti-TAP Ab (Thermo Fisher Scientific, CAB1001) was added and incubated for 4 hours on a rotator wheel at 4°C. To this extract, 100 μl bed volume of pre-equilibrated Protein A^PLUS^ Agarose Beads was added and incubated at 4°C on the rotator wheel overnight. The beads are washed thrice with ten volumes of Buffer A by rotating on the rotator wheel at 4°C for ten minutes each. Finally, elution is performed by boiling the beads in 40ul of SDS loading dye for 5 minutes. Samples are now ready to be analyzed on SDS-PAGE gel followed by Western blot analysis.

### Protein analyses by western blot

Total protein was isolated from specific yeast strains (TAP-Rrp47) grown overnight at 30°C in YPD broth. Following centrifugation at 5000 rpm for 7 min, the cell pellets were quickly frozen in liquid nitrogen and stored at −70°C. Frozen pellets were thawed on ice and resuspended in 1 ml of Buffer A (50 mM Tris–HCl pH 7.5, 150 mM NaCl, 5 mM EDTA, 1 mM DTT, 1 mM PMSF) supplemented with Protease Inhibitor Cocktail (Abcam ab201111, UK) and the cells were lysed by vortexing 10–15 times with glass beads followed by clarification of the particulate fraction by centrifugation. Supernatants were collected by centrifugation at 11 000 rpm for 20 min and saved as the total soluble protein fraction for further analysis. Protein concentration was determined by Bradford reagent assay kit (Bio-Rad Inc., Valencia, CA, USA). For Western Analysis, 60 μg of total protein was used, which was resolved in 10% SDS polyacrylamide gel. The separated proteins were transferred to PVDF membrane at 50-100mA O/N for immunoblotting. Blots were blocked with 5% skimmed milk in Tris-buffered saline (10mMTris, 150mMNaCl, 0.1% Tween 20) and incubated with primary antibodies for specific proteins 1 h at room temperature diluted in 5% BSA (Table S5). Blots were further washed in 1X TBS with 0.1% Tween 20 (TBST) and incubated in HRP-conjugated secondary anti-rabbit or anti-mouse antibody, each diluted at 1:3000 in wash buffer for 1 h at room temperature. Immunoreactive bands were developed and detected by chemiluminescence (ECL imager kit, Abcam), and the images were captured by X-ray film or myECL Chemidoc Imager (Thermo Scientific, USA).

### Protein Purification using Gel Filtration Column Chromatography

Yeast cells were grown O/N at 30 ° C in 200-250 ml YPD broth until O.D 600 =3, followed by cell harvesting, followed by Protein extraction using the standard protocol mentioned before. The protein extract was ultra-centrifuged twice at 40000 g for 10 minutes. The final sup was collected and stored at −80 degrees ultra freezer for future use. To perform a separation, gel filtration bead (P-200 beads from BioRad) was soaked in water O/N at room temp. It was then packed into a column to form a packed bed. The medium is a porous matrix of spherical particles with chemical and physical stability and inertness (lack of reactivity and adsorptive properties). The packed bed was equilibrated with Tris-NaCl buffer, which fills the pores of the matrix and the space between the particles. The liquid inside the pores, or stationary phase, is in equilibrium with the liquid outside the particles, or mobile phase. The protein extract was then run on the column, and 100-500 μl of samples were eluted isocratically, so there is no need to use different buffers during the separation. However, a wash step using the running buffer is usually included at the end of separation to remove molecules that may have been retained on the column and prepare the column for a new run. Finally, the eluates are used for the SDS-PAGE and Western Blotting analysis. (The proteins eluates should be precipitated to get the purified concentrated protein and dissolved in pure Milli-Q water before any high-end experimental analysis).

### Genomic Data Mining and Analysis (RNA Sequencing)

We performed a re-analysis of a previously submitted dataset (117) available from Gene Expression Omnibus (GEO) under the accession number GSE135056. The entire analysis was done using the public server (usegalaxy.org) of The Galaxy Project (118). The FASTQ files for the WT and *rrp6-*Δ RNA-Seq data were imported into the galaxy server using the SRA import tool **Faster Download and Extract Reads in FASTQ** format from NCBI SRA (119). Quality checking and trimming as required was done with FastQC (120), Cutadapt (121), and MultiQC (122).

The processed reads were then aligned to the **R64-1-1 (GCA_000146045.2)** reference genome from ensemble.org with **RNA STAR** (123). The annotation file used as a gene model was **R64-1-1.101.gtf**, also downloaded from ensemble.org. To confirm that most of the reads have aligned to exons rather than intergenic regions (due to DNA contamination), we used **Read Distribution** from the **RSeQC** (124) tool suite to check the percentage of reads mapping to known genomic features. **featureCounts** (125) was then used to count the number of reads per annotated gene. To use featureCounts, we needed to know if the sequence library was stranded or not. To predict the strandedness of the library, **Infer Experiment** from the **RSeQC** (124) tool suite was used, and the instructions were followed as is, from the galaxy tutorial, **Reference-based RNA-Seq data analysis** (126)(127) to determine how the sequencing library was configured. The information thus obtained from Infer Experiment (124) was used in featureCounts (125) with the option “Count fragments instead of reads” enabled to ultimately obtain the counts files for all the samples.

The counts files for the technical replicates were merged by summing up the counts for each feature before feeding them into **edgeR** (101–103) so that the statistical analysis is performed on the biological replicates only so as to obtain a better picture about the variance among the biological replicates. Since the number of replicates per condition was less than 12, the tools of choice (128) were either edgeR (101–103) or DESeq2 (129). Coverage Plots were created using Integrative Genomics Viewer (130). **Volcano Plot** and **heatmap2** tools were used to create the volcano plots and heatmaps.

### Statistical analyses

The quantitative experiments reported in this paper (RNA steady-state levels, transcription shut-off Decay experiments, and co-localization index) were performed using at least three independent sample size (biological replicate, *N* = 3 to 5).

A given yeast strain was grown and treated under the same experimental conditions independently before a given experiment was conducted for each biological replicate. In technical replication, repetition/analyses of the same biological replicate sample were performed many times. To establish the variability (experimental error) involved in the analysis, a technical replicate was used, which allows the experimenter to set the confidence limits for what are significant data. All the statistical parameters such as mean, standard deviations(SD), Standard Error of Mean(SEM), median, and P values were calculated using OriginPro8 and GraphPad Prism version 8.4 (GraphPad Software Inc., San Diego, CA, USA). P-values were calculated using Student’s two-tailed t-test (unpaired) using the same program. The values of different statistical parameters (Biological Replicate, N, mean, median, SD, SEM, and P-value) corresponding to the individual experiments are given in a separate MS Excel File.

## Acknowledgments

We gratefully acknowledge Prof. Scott Butler (Department of Microbiology and Immunology, University of Rochester) and Dr. Satarupa Das (Jadavpur University, India), and members of Das laboratory for critically reading the manuscript and useful comments and suggestions. The authors gratefully acknowledge Prof. Scott Butler (Department of Microbiology and Immunology and Microbiology, University of Rochester), Prof. Krishnaveni Mishra (School of Life Science, University of Hyderabad), and Prof. Rachid Rahmouni (French National Centre for Scientific Research, France) for sharing many useful the plasmids and yeast strains. We also thank the anonymous reviewers for their critical comments and constructive suggestions, which enriched this manuscript.

## 4. Author contributions

AC and SD performed experiments, AC and BD designed the experiments and analyzed the data, MB analyzed the RNA-Seq data, AC and BD drafted and wrote the manuscript.

## References

1. Venema,J. and Tollervey,D. (1999) Ribosome Synthesis in Saccharomyces cerevisiae. Annu. Rev. Genet., 33, 261–311.

2. Woolford,J.L. and Baserga,S.J. (2013) Ribosome biogenesis in the yeast Saccharomyces cerevisiae. Genetics, 195, 643–681.

3. Henras,A.K., Plisson-Chastang,C., O’Donohue,M.F., Chakraborty,A. and Gleizes,P.E. (2015) An overview of pre-ribosomal RNA processing in eukaryotes. Wiley Interdiscip. Rev. RNA, 6, 225–242.

4. Wikipedia,F. (2013) COPB2. EMBO J., 2, 4–7.

5. Elela,S.A., Igel,H. and Ares,M. (1996) RNase III cleaves eukaryotic preribosomal RNA at a U3 snoRNP-dependent site. Cell, 85, 115–124.

6. Kempers-Veenstra,A.E., Oliemans,J., Offenberg,H., Dekker,A.F., Piper,P.W., Planta,R.J. and Klootwijk,J. (1986) 3’-End formation of transcripts from the yeast rRNA operon. EMBO J., 5, 2703–2710.

7. Kufel,J., Dichtl,B. and Tollervey,D. (1999) Yeast Rnt1p is required for cleavage of the pre-ribosomal RNA in the 3’ ETS but not the 5’ ETS. RNA, 5, 909–917.

8. Oeffinger,M., Zenklusen,D., Ferguson,A., Wei,K.E., El Hage,A., Tollervey,D., Chait,B.T., Singer,R.H. and Rout,M.P. (2009) Rrp17p Is a Eukaryotic Exonuclease Required for 5’ End Processing of Pre-60S Ribosomal RNA. Mol. Cell, 36, 768–781.

9. Amberg,D.C., Goldstein,A.L. and Cole,C.N. (1992) Isolation and characterization of RAT1: an essential gene of Saccharomyces cerevisiae required for the efficient nucleocytoplasmic trafficking of mRNA. Genes Dev, 6, 1173–1189.

10. Henry,Y., Wood,H., Morrissey,J.P., Petfalski,E., Kearsey,S. and Tollervey,D. (1994) The 5’ end of yeast 5.8S rRNA is generated by exonucleases from an upstream cleavage site. EMBO J., 13, 2452–2463.

11. Kim,M., Krogan,N.J., Vasiljeva,L., Rando,O.J., Nedea,E., Greenblatt,J.F. and Buratowski,S. (2004) The yeast Rat1 exonuclease promotes transcription termination by RNA polymerase II. Nature, 432, 517–522.

12. Fang,F., Phillips,S. and Butler,J.S. (2005) Rat1p and Rai1p function with the nuclear exosome in the processing and degradation of rRNA precursors. RNA, 11, 1571–8.

13. Axt,K., French,S.L., Beyer,A.L. and Tollervey,D. (2014) Kinetic analysis demonstrates a requirement for the Rat1 exonuclease in cotranscriptional pre-rRNA cleavage. PLoS One, 9.

14. Thomson,E. and Tollervey,D. (2010) The Final Step in 5.8S rRNA Processing Is Cytoplasmic in Saccharomyces cerevisiae. Mol. Cell. Biol., 30, 976–984.

15. Allmang,C., Kufel,J., Chanfreau,G., Mitchell,P., Petfalski,E. and Tollervey,D. (1999) Functions of the exosome in rRNA, snoRNA and snRNA synthesis. EMBO J, 18, 5399–5410.

16. Allmang,C. and Tollervey,D. (1998) The role of the 3’ external transcribed spacer in yeast pre-rRNA processing. J. Mol. Biol., 278, 67–78.

17. de la Cruz,J., Kressler,D., Tollervey,D. and Linder,P. (1998) Dob1p (Mtr4p) is a putative ATP-dependent RNA helicase required for the 3’ end formation of 5.8S rRNA in Saccharomyces cerevisiae. EMBO J, 17, 1128–1140.

18. Briggs,M.W., Burkard,K.T. and Butler,J.S. (1998) Rrp6p, the yeast homologue of the human PM-Scl 100-kDa autoantigen, is essential for efficient 5.8 S rRNA 3’ end formation. J Biol Chem, 273, 13255–13263.

19. Houseley,J. and Tollervey,D. (2006) Yeast Trf5p is a nuclear poly(A) polymerase. EMBO Rep, 7, 205–211.

20. Piper,P.W., Bellatin,J.A. and Lockheart,A. (1983) Altered maturation of sequences at the 3’ terminus of 5S gene transcripts in a Saccharomyces cerevisiae mutant that lacks a RNA processing endonuclease. EMBO J., 2, 353–359.

21. Kufel,J. and Grzechnik,P. (2019) Small Nucleolar RNAs Tell a Different Tale. Trends Genet., 35, 104–117.

22. Dieci,G., Preti,M. and Montanini,B. (2009) Eukaryotic snoRNAs: A paradigm for gene expression flexibility. Genomics, 94, 83–88.

23. Huang,Y. and Carmichael,G.G. (1996) Role of polyadenylation in nucleocytoplasmic transport of mRNA. Mol. Cell. Biol., 16, 1534–1542.

24. Dunn,E.F., Hammell,C.M., Hodge,C.A. and Cole,C.N. (2005) Yeast poly(A)-binding protein, Pab1, and PAN, a poly(A) nuclease complex recruited by Pab1, connect mRNA biogenesis to export. Genes Dev., 19, 90–103.

25. Proudfoot,N.J. (2016) Transcriptional termination in mammals: Stopping the RNA polymerase II juggernaut. Science (80-.)., 352.

26. Porrua,O., Boudvillain,M. and Libri,D. (2016) Transcription Termination: Variations on Common Themes. Trends Genet., 32, 508–522.

27. Porrua,O. and Libri,D. (2013) A bacterial-like mechanism for transcription termination by the Sen1p helicase in budding yeast. Nat. Struct. Mol. Biol., 20, 884–891.

28. Lemay,J.F., Marguerat,S., Larochelle,M., Liu,X., van Nues,R., Hunyadkürti,J., Hoque,M., Tian,B., Granneman,S., Bähler,J., et al. (2016) The Nrd1-like protein Seb1 coordinates cotranscriptional 3’ end processing and polyadenylation site selection. Genes Dev., 30, 1558–1572.

29. Wittmann,S., Renner,M., Watts,B.R., Adams,O., Huseyin,M., Baejen,C., El Omari,K., Kilchert,C., Heo,D.H., Kecman,T., et al. (2017) The conserved protein Seb1 drives transcription termination by binding RNA polymerase II and nascent RNA. Nat. Commun., 8.

30. LaCava,J., Houseley,J., Saveanu,C., Petfalski,E., Thompson,E., Jacquier,A. and Tollervey,D. (2005) RNA degradation by the exosome is promoted by a nuclear polyadenylation complex. Cell, 121, 713–724.

31. Schmidt,K. and Butler,J.S. (2013) Nuclear RNA surveillance: role of TRAMP in controlling exosome specificity. Wiley Interdiscip Rev RNA, 4, 217–231.

32. Grzechnik,P. and Kufel,J. (2008) Polyadenylation Linked to Transcription Termination Directs the Processing of snoRNA Precursors in Yeast. Mol. Cell, 32, 247–258.

33. Ooi,A.T., Gower,A.C., Zhang,K.X., Vick,J.L., Hong,L., Nagao,B., Wallace,W.D., Elashoff,D.A., Walser,T.C., Dubinett,S.M., et al. (2014) Molecular profiling of premalignant lesions in lung squamous cell carcinomas identifies mechanisms involved in stepwise carcinogenesis. Cancer Prev Res, 7, 487–495.

34. Ghazal,G., Ge,D., Gervais-Bird,J., Gagnon,J. and Abou Elela,S. (2005) Genome-Wide Prediction and Analysis of Yeast RNase III-Dependent snoRNA Processing Signals. Mol. Cell. Biol., 25, 2981–2994.

35. Villa,T., Ceradini,F., Presutti,C. and Bozzoni,I. (1998) Processing of the Intron-Encoded U18 Small Nucleolar RNA in the Yeast Saccharomyces cerevisiaeRelies on Both Exo- and Endonucleolytic Activities. Mol. Cell. Biol., 18, 3376–3383.

36. Wlotzka,W., Kudla,G., Granneman,S. and Tollervey,D. (2011) The nuclear RNA polymerase II surveillance system targets polymerase III transcripts. EMBO J, 30, 1790–1803.

37. Siliciano,P.G., Jones,M.H. and Guthrie,C. (1987) Saccharomyces cerevisiae has a Ul-like small nuclear RNA with unexpected properties. Science (80-.)., 237, 1484–1487.

38. Kiss,T. (2004) Biogenesis of small nuclear RNPs. J. Cell Sci., 117, 5949–5951.

39. Liu,Q., Greimann,J.C. and Lima,C.D. (2006) Reconstitution, activities, and structure of the eukaryotic RNA exosome. Cell, 127, 1223–1237.

40. Dziembowski,A., Lorentzen,E., Conti,E. and Seraphin,B. (2007) A single subunit, Dis3, is essentially responsible for yeast exosome core activity. Nat Struct Mol Biol, 14, 15–22.

41. Mitchell,P., Petfalski,E., Shevchenko,A., Mann,M. and Tollervey,D. (1997) The exosome: A conserved eukaryotic RNA processing complex containing multiple 3’→5’ exoribonucleases. Cell, 91, 457–466.

42. Burkard,K.T. and Butler,J.S. (2000) A nuclear 3’-5’ exonuclease involved in mRNA degradation interacts with Poly(A) polymerase and the hnRNA protein Npl3p. Mol Cell Biol, 20, 604–616.

43. Houseley,J. and Tollervey,D. (2009) The Many Pathways of RNA Degradation. Cell, 136, 763–776.

44. Vanacova,S., Wolf,J., Martin,G., Blank,D., Dettwiler,S., Friedlein,A., Langen,H., Keith,G. and Keller,W. (2005) A new yeast poly(A) polymerase complex involved in RNA quality control. PLoS Biol, 3, e189.

45. Wyers,F., Rougemaille,M., Badis,G., Rousselle,J.C., Dufour,M.E., Boulay,J., Regnault,B., Devaux,F., Namane,A., Seraphin,B., et al. (2005) Cryptic pol II transcripts are degraded by a nuclear quality control pathway involving a new poly(A) polymerase. Cell, 121, 725–737.

46. Maity,A., Chaudhuri,A. and Das,B. (2016) DRN and TRAMP degrade specific and overlapping aberrant mRNAs formed at various stages of mRNP biogenesis in Saccharomyces cerevisiae. FEMS Yeast Res., 16.

47. Das,B., Guo,Z., Russo,P., Chartrand,P. and Sherman,F. (2000) The role of nuclear cap binding protein Cbc1p of yeast in mRNA termination and degradation. Mol. Cell. Biol., 20.

48. Das,B., Butler,J.S. and Sherman,F. (2003) Degradation of normal mRNA in the nucleus of Saccharomyces cerevisiae. Mol. Cell. Biol., 23.

49. Das,S., Saha,U. and Das,B. (2014) Cbc2p, Upf3p and eIF4G are components of the DRN (Degradation of mRNA in the Nucleus) in Saccharomyces cerevisiae. FEMS Yeast Res., 14.

50. Cloutier,S.C., Ma,W.K., Nguyen,L.T. and Tran,E.J. (2012) The DEAD-box RNA helicase Dbp2 connects RNA quality control with repression of aberrant transcription. J. Biol. Chem., 287, 26155–26166.

51. Eichinger,C.S. and Jentsch,S. (2010) Synaptonemal complex formation and meiotic checkpoint signaling are linked to the lateral element protein red1. Proc. Natl. Acad. Sci. U. S. A., 107, 11370–11375.

52. Allmang,C., Petfalski,E., Podtelejnikov,A., Mann,M., Tollervey,D. and Mitchell,P. (1999) The yeast exosome and human PM-Scl are related complexes of 3’ → 5’ exonucleases. Genes Dev., 13, 2148–2158.

53. Allmang,C., Mitchell,P., Petfalski,E. and Tollervey,D. (2000) Degradation of ribosomal RNA precursors by the exosome. Nucleic Acids Res, 28, 1684–1691.

54. Chen,C.Y., Gherzi,R., Ong,S.E., Chan,E.L., Raijmakers,R., Pruijn,G.J., Stoecklin,G., Moroni,C., Mann,M. and Karin,M. (2001) AU binding proteins recruit the exosome to degrade ARE-containing mRNAs. Cell, 107, 451–464.

55. Brouwer,R., Vree Egberts,W.T.M., Hengstman,G.J.D., Raijmakers,R., van Engelen,B.G.M., Seelig,H.P., Renz,M., Mierau,R., Genth,E., Pruijn,G.J.M., et al. (2002) Autoantibodies directed to novel components of the PM/Scl complex, the human exosome. Arthritis Res., 4, 134–8.

56. Butler,J.S. (2002) The yin and yang of the exosome. Trends Cell Biol, 12, 90–96.

57. Vanacova,S. and Stefl,R. (2007) The exosome and RNA quality control in the nucleus. EMBO Rep, 8, 651–657.

58. Brouwer,R., Pruijn,G.J.M. and van Venrooij,W.J. (2001) The human exosome: An autoantigenic complex of exoribonucleases in myositis and scleroderma. Arthritis Res., 3, 102–106.

59. Estévez,A.M., Lehner,B., Sanderson,C.M., Ruppert,T. and Clayton,C. (2003) The roles of intersubunit interactions in exosome stability. J. Biol. Chem., 278, 34943–34951.

60. Estévez A M, Kempf,T. and Clayton C (2001) The Exosome of Trypanosoma Brucei. EMBO J., 20, 3831–3839.

61. Andrulis,E.D., Werner,J., Nazarian,A., Erdjument-Bromage,H., Tempst,P. and Lis,J.T. (2002) The RNA processing exosome is linked to elongating RNA polymerase II in Drosophila. Nature, 420, 837–841.

62. Evguenieva-Hackenberg,E., Walter,P., Hochleitner,E., Lottspeich,F. and Klug,G. (2003) An exosome-like complex in Sulfolobus solfataricus. EMBO Rep., 4, 889–893.

63. Büttner,K., Wenig,K. and Hopfner,K.P. (2005) Structural framework for the mechanism of archaeal exosomes in RNA processing. Mol. Cell, 20, 461–471.

64. Yang,C.C., Wang,Y.T., Hsiao,Y.Y., Doudeva,L.G., Kuo,P.H., Chow,S.Y. and Yuan,H.S. (2010) Structural and biochemical characterization of CRN-5 and Rrp46: An exosome component participating in apoptotic DNA degradation. RNA, 16, 1748–1759.

65. Kiss,D.L. and Andrulis,E.D. (2011) The exozyme model: a continuum of functionally distinct complexes. RNA, 17, 1–13.

66. Kadaba,S., Krueger,A., Trice,T., Krecic,A.M., Hinnebusch,A.G. and Anderson,J. (2004) Nuclear surveillance and degradation of hypomodified initiator tRNA Met in S. cerevisiae. Genes Dev., 18, 1227–1240.

67. Houalla,R., Devaux,F., Fatica,A., Kufel,J., Barrass,D., Torchet,C. and Tollervey,D. (2006) Microarray detection of novel nuclear RNA substrates for the exosome. Yeast, 23, 439–454.

68. Callahan,K.P. and Butler,J.S. (2008) Evidence for core exosome independent function of the nuclear exoribonuclease Rrp6p. Nucleic Acids Res., 36, 6645–55.

69. Schneider,C., Anderson,J.T. and Tollervey,D. (2007) The Exosome Subunit Rrp44 Plays a Direct Role in RNA Substrate Recognition. Mol. Cell, 27, 324–331.

70. Graham,A.C., Kiss,D.L. and Andrulis,E.D. (2009) Core exosome - Independent roles for Rrp6 in cell cycle progression. Mol. Biol. Cell, 20, 2242–2253.

71. Kiss,D.L. and Andrulis,E.D. (2010) Genome-wide analysis reveals distinct substrate specificities of Rrp6, Dis3, and core exosome subunits. RNA, 16, 781–791.

72. Slomovic,S., Fremder,E., Staals,R.H.G., Pruijn,G.J.M. and Schuster,G. (2010) Addition of poly(A) and poly(A)-rich tails during RNA degradation in the cytoplasm of human cells. Proc. Natl. Acad. Sci. U. S. A., 107, 7407–7412.

73. Gavin,A.C., Bösche,M., Krause,R., Grandi,P., Marzioch,M., Bauer,A., Schultz,J., Rick,J.M., Michon,A.M., Cruciat,C.M., et al. (2002) Functional organization of the yeast proteome by systematic analysis of protein complexes. Nature, 415, 141–147.

74. Krogan,N.J., Peng,W.T., Cagney,G., Robinson,M.D., Haw,R., Zhong,G., Guo,X., Zhang,X., Canadien,V., Richards,D.P., et al. (2004) High-Definition Macromolecular Composition of Yeast RNA-Processing Complexes. Mol. Cell, 13, 225–239.

75. Krogan,N.J., Cagney,G., Yu,H., Zhong,G., Guo,X., Ignatchenko,A., Li,J., Pu,S., Datta,N., Tikuisis,A.P., et al. (2006) Global landscape of protein complexes in the yeast Saccharomyces cerevisiae. Nature, 440, 637–643.

76. Synowsky,S.A., van Wijk,M., Raijmakers,R. and Heck,A.J.R. (2009) Comparative Multiplexed Mass Spectrometric Analyses of Endogenously Expressed Yeast Nuclear and Cytoplasmic Exosomes. J. Mol. Biol., 385, 1300–1313.

77. Jostins,L., Ripke,S., Weersma,R.K., Duerr,R.H., McGovern,D.P., Hui,K.Y., Lee,J.C., Schumm,L.P., Sharma,Y., Anderson,C.A., et al. (2012) Host-microbe interactions have shaped the genetic architecture of inflammatory bowel disease. Nature, 491, 119–124.

78. Graham,A.C., Kiss,D.L. and Andrulis,E.D. (2006) Differential distribution of exosome subunits at the nuclear lamina and in cytoplasmic foci. Mol. Biol. Cell, 17, 1399–1409.

79. Callahan,K.P. and Butler,J.S. (2010) TRAMP complex enhances RNA degradation by the nuclear exosome component Rrp6. J Biol Chem, 285, 3540–3547.

80. Wyers,F., Rougemaille,M., Badis,G., Rousselle,J.C., Dufour,M.E., Boulay,J., Régnault,B., Devaux,F., Namane,A., Séraphin,B., et al. (2005) Cryptic Pol II transcripts are degraded by a nuclear quality control pathway involving a new poly(A) polymerase. Cell, 121, 725–737.

81. Chekanova,J.A., Gregory,B.D., Reverdatto,S. V, Chen,H., Kumar,R., Hooker,T., Yazaki,J., Li,P., Skiba,N., Peng,Q., et al. (2007) Genome-wide high-resolution mapping of exosome substrates reveals hidden features in the Arabidopsis transcriptome. Cell, 131, 1340–1353.

82. Evguenieva-Hackenberg,E., Roppelt,V., Finsterseifer,P. and Klug,G. (2008) Rrp4 and Csl4 are needed for efficient degradation but not for polyadenylation of synthetic and natural RNA by the archaeal exosome. Biochemistry, 47, 13158–13168.

83. Roppelt,V., Klug,G. and Evguenieva-Hackenberg,E. (2010) The evolutionarily conserved subunits Rrp4 and Csl4 confer different substrate specificities to the archaeal exosome. FEBS Lett., 584, 2931–2936.

84. Kuai,L., Fang,F., Butler,J.S. and Sherman,F. (2004) Polyadenylation of rRNA in Saccharomyces cerevisiae. Proc. Natl. Acad. Sci. U. S. A., 101, 8581–8586.

85. Briggs,M.W., Burkard,K.T.D. and Butler,J.S. (1998) Rrp6p, the yeast homologue of the human PM-Scl 100-kDa autoantigen, is essential for efficient 5.8 S rRNA 3’ end formation. J. Biol. Chem., 273, 13255–13263.

86. Butler,J.S. and Mitchell,P. (2011) Rrp6, rrp47 and cofactors of the nuclear exosome. Adv Exp Med Biol, 702, 91–104.

87. Tudek,A., Porrua,O., Kabzinski,T., Lidschreiber,M., Kubicek,K., Fortova,A., Lacroute,F., Vanacova,S., Cramer,P., Stefl,R., et al. (2014) Molecular basis for coordinating transcription termination with noncoding RNA degradation. Mol Cell, 55, 467–481.

88. Vasiljeva,L. and Buratowski,S. (2006) Nrd1 interacts with the nuclear exosome for 3’ processing of RNA polymerase II transcripts. Mol Cell, 21, 239–248.

89. Chanfreau,G., Legrain,P. and Jacquier,A. (1998) Yeast RNase III as a key processing enzyme in small nucleolar RNAs metabolism. J. Mol. Biol., 284, 975–988.

90. Kufel,J., Allmang,C., Chanfreau,G., Petfalski,E., Lafontaine,D.L.J. and Tollervey,D. (2000) Precursors to the U3 Small Nucleolar RNA Lack Small Nucleolar RNP Proteins but Are Stabilized by La Binding. Mol. Cell. Biol., 20, 5415–5424.

91. Feigenbutz,M., Jones,R., Besong,T.M., Harding,S.E. and Mitchell,P. (2013) Assembly of the yeast exoribonuclease Rrp6 with its associated cofactor Rrp47 occurs in the nucleus and is critical for the controlled expression of Rrp47. J Biol Chem, 288, 15959–15970.

92. Milligan,L., Decourty,L., Saveanu,C., Rappsilber,J., Ceulemans,H., Jacquier,A. and Tollervey,D. (2008) A yeast exosome cofactor, Mpp6, functions in RNA surveillance and in the degradation of noncoding RNA transcripts. Mol Cell Biol, 28, 5446–5457.

93. Stead,J.A., Costello,J.L., Livingstone,M.J. and Mitchell,P. (2007) The PMC2NT domain of the catalytic exosome subunit Rrp6p provides the interface for binding with its cofactor Rrp47p, a nucleic acid-binding protein. Nucleic Acids Res, 35, 5556–5567.

94. Mitchell,P., Petfalski,E., Houalla,R., Podtelejnikov,A., Mann,M. and Tollervey,D. (2003) Rrp47p is an exosome-associated protein required for the 3’ processing of stable RNAs. Mol Cell Biol, 23, 6982–6992.

95. Stuparevic,I., Mosrin-Huaman,C., Hervouet-Coste,N., Remenaric,M. and Rahmouni,A.R. (2013) Cotranscriptional recruitment of RNA exosome cofactors Rrp47p and Mpp6p and two distinct Trf-Air-Mtr4 polyadenylation (TRAMP) complexes assists the exonuclease Rrp6p in the targeting and degradation of an aberrant messenger ribonucleoprotein particle (mRN. J. Biol. Chem., 288, 31816–31829.

96. Phillips,S. and Butler,J.S. (2003) Contribution of domain structure to the RNA 3’ end processing and degradation functions of the nuclear exosome subunit Rrp6p. RNA, 9, 1098–1107.

97. Lingner,J., Kellermann,J. and Keller,W. (1991) Cloning and expression of the essential gene for poly(A) polymerase from S. cerevisiae. Nature, 354, 496–498.

98. Patel,D. and Butler,J.S. (1992) Conditional defect in mRNA 3’ end processing caused by a mutation in the gene for poly(A) polymerase. Mol. Cell. Biol., 12, 3297–3304.

99. Vaňáčová,Š., Wolf,J., Martin,G., Blank,D., Dettwiler,S., Friedlein,A., Langen,H., Keith,G. and Keller,W. (2005) A new yeast poly(A) polymerase complex involved in RNA quality control. PLoS Biol., 3, 0986–0997.

100. Wang,E.T., Taliaferro,J.M., Lee,J.A., Sudhakaran,I.P., Rossoll,W., Gross,C., Moss,K.R. and Bassell,G.J. (2016) Dysregulation of mRNA localization and translation in genetic disease. J. Neurosci., 36, 11418–11426.

101. Robinson,M.D., McCarthy,D.J. and Smyth,G.K. (2010) edgeR: a Bioconductor package for differential expression analysis of digital gene expression data. Bioinformatics, 26, 139–140.

102. McCarthy,D.J., Chen,Y. and Smyth,G.K. (2012) Differential expression analysis of multifactor RNA-Seq experiments with respect to biological variation. Nucleic Acids Res., 40, 4288–4297.

103. Liu,R., Holik,A.Z., Su,S., Jansz,N., Chen,K., Leong,H.S., Blewitt,M.E., Asselin-Labat,M.-L., Smyth,G.K. and Ritchie,M.E. (2015) Why weight? Modelling sample and observational level variability improves power in RNA-seq analyses. Nucleic Acids Res., 43, e97–e97.

104. Petfalski,E., Dandekar,T., Henry,Y. and Tollervey,D. (1998) Processing of the Precursors to Small Nucleolar RNAs and rRNAs Requires Common Components. Mol. Cell. Biol., 18, 1181–1189.

105. van Hoof,A., Staples,R.R., Baker,R.E. and Parker,R. (2000) Function of the ski4p (Csl4p) and Ski7p proteins in 3’-to-5’ degradation of mRNA. Mol Cell Biol, 20, 8230–8243.

106. Heo,D.H., Yoo,I., Kong,J., Lidschreiber,M., Mayer,A., Choi,B.Y., Hahn,Y., Cramer,P., Buratowski,S. and Kim,M. (2013) The RNA polymerase 2 C-terminal domain-interacting domain of yeast Nrd1 contributes to the choice of termination pathway and couples to RNA processing by the nuclear exosome. J. Biol. Chem., 288, 36676–36690.

107. Fox,M.J. and Mosley,A.L. (2016) Rrp6: Integrated roles in nuclear RNA metabolism and transcription termination. Wiley Interdiscip. Rev. RNA, 7, 91–104.

108. Gudipati,R.K., Xu,Z., Lebreton,A., Seraphin,B., Steinmetz,L.M., Jacquier,A. and Libri,D. (2012) Extensive degradation of RNA precursors by the exosome in wild-type cells. Mol Cell, 48, 409–421.

109. Feigenbutz,M., Garland,W., Turner,M. and Mitchell,P. (2013) The exosome cofactor Rrp47 is critical for the stability and normal expression of its associated exoribonuclease Rrp6 in Saccharomyces cerevisiae. PLoS One, 8, e80752.

110. Kim,M., Vasiljeva,L., Rando,O.J., Zhelkovsky,A., Moore,C. and Buratowski,S. (2006) Distinct pathways for snoRNA and mRNA termination. Mol Cell, 24, 723–734.

111. Fox,M.J., Gao,H., Smith-Kinnaman,W.R., Liu,Y. and Mosley,A.L. (2015) The Exosome Component Rrp6 Is Required for RNA Polymerase II Termination at Specific Targets of the Nrd1-Nab3 Pathway. PLoS Genet., 11, 1–26.

112. Carneiro,T., Carvalho,C., Braga,J., Rino,J., Milligan,L., Tollervey,D. and Carmo-Fonseca,M. (2007) Depletion of the Yeast Nuclear Exosome Subunit Rrp6 Results in Accumulation of Polyadenylated RNAs in a Discrete Domain within the Nucleolus. Mol. Cell. Biol., 27, 4157–4165.

113. Albert,B., Kos-Braun,I.C., Henras,A.K., Dez,C., Rueda,M.P., Zhang,X., Gadal,O., Kos,M. and Shore,D. (2019) A ribosome assembly stress response regulates transcription to maintain proteome homeostasis. Elife, 8.

114. Szaflarski,W., Sowiński,M., Leśniczak,M., Ojha,S., Aulas,A., Dave,D., Malla,S., Anderson,P., Ivanov,P. and Lyons,S.M. (2020) A novel stress response pathway regulates rRNA biogenesis. bioRxiv, 10.1101/2020.08.16.250183.

115. Sherman,F. (1991) Getting Started with Yeast. Methods Enzymol., 194, 3–21.

116. Molecular Cloning: A Laboratory Manual (Fourth Edition).

117. Victorino,J.F., Fox,M.J., Smith-Kinnaman,W.R., Peck Justice,S.A., Burriss,K.H., Boyd,A.K., Zimmerly,M.A., Chan,R.R., Hunter,G.O., Liu,Y., et al. (2020) RNA Polymerase II CTD phosphatase Rtr1 fine-tunes transcription termination. PLOS Genet., 16, e1008317.

118. Afgan,E., Baker,D., Batut,B., van den Beek,M., Bouvier,D., Čech,M., Chilton,J., Clements,D., Coraor,N., Grüning,B.A., et al. (2018) The Galaxy platform for accessible, reproducible and collaborative biomedical analyses: 2018 update. Nucleic Acids Res., 46, W537–W544.

119. Leinonen,R., Sugawara,H., Shumway,M. and Collaboration,I.N.S.D. (2011) The sequence read archive. Nucleic Acids Res., 39, D19–D21.

120. Andrews,S. FastQC A Quality Control tool for High Throughput Sequence Data.

121. Martin,M. (2011) Cutadapt removes adapter sequences from high-throughput sequencing reads. EMBnet.journal; Vol 17, No 1 Next Gener. Seq. Data Anal. - 10.14806/ej.17.1.200.

122. Ewels,P., Magnusson,M., Lundin,S. and Käller,M. (2016) MultiQC: summarize analysis results for multiple tools and samples in a single report. Bioinformatics, 32, 3047–3048.

123. Dobin,A., Davis,C.A., Schlesinger,F., Drenkow,J., Zaleski,C., Jha,S., Batut,P., Chaisson,M. and Gingeras,T.R. (2013) STAR: ultrafast universal RNA-seq aligner. Bioinformatics, 29, 15–21.

124. Wang,L., Wang,S. and Li,W. (2012) RSeQC: quality control of RNA-seq experiments. Bioinformatics, 28, 2184–2185.

125. Liao,Y., Smyth,G.K. and Shi,W. (2014) featureCounts: an efficient general purpose program for assigning sequence reads to genomic features. Bioinformatics, 30, 923–930.

126. Batut,B., Freeberg,M., Heydarian,M., Erxleben,A., Videm,P., Blank,C., Doyle,M., Soranzo,N. and van Heusden,P. (2020) Reference-based RNA-Seq data analysis (Galaxy Training Materials).

127. Batut,B., Hiltemann,S., Bagnacani,A., Baker,D., Bhardwaj,V., Blank,C., Bretaudeau,A., Brillet-Guéguen,L., Čech,M., Chilton,J., et al. (2018) Community-Driven Data Analysis Training for Biology. Cell Syst., 6, 752--758.e1.

128. Schurch,N.J., Schofield,P., Gierliński,M., Cole,C., Sherstnev,A., Singh,V., Wrobel,N., Gharbi,K., Simpson,G.G., Owen-Hughes,T., et al. (2016) How many biological replicates are needed in an RNA-seq experiment and which differential expression tool should you use? RNA, 22, 839–851.

129. Love,M.I., Huber,W. and Anders,S. (2014) Moderated estimation of fold change and dispersion for RNA-seq data with DESeq2. Genome Biol., 15, 550.

130. Robinson,J.T., Thorvaldsdóttir,H., Winckler,W., Guttman,M., Lander,E.S., Getz,G. and Mesirov,J.P. (2011) Integrative genomics viewer. Nat. Biotechnol., 29, 24–26.

