## Supplementary Tables S1-5 and Figure S1-3 for "Rrp6p/Rrp47p constitutes an independent nuclear turnover system of mature small non-coding RNAs in *Saccharomyces cerevisiae*"

**RESEARCH ARTICLE** **Submitted to *Nature Communications***

Submitted December 7, 2020

### SUPPLEMENTARY DATA

***Table S1: List and Genotypes of Yeast Strains used in this study***

| **Strain No.** | **Complete genotype** | **Abbreviated Genotype** | **Reference** |
|---|---|---|---|
| yBD 263 | MATa cyc1-512 ura3-52 trp2-1 *his3*::hisG | WT | *This Work |
| yBD 264 | MATa cyc1-512 ura3-52 trp2-1 *his3::hisG* cbc1::URA3 | cbc1Δ | *This Work |
| yBD 265 | MATa cyc1-512 ura3-52 trp2-1 *his3::hisG* rrp6::URA3 | rrp6Δ | *This Work |
| yBD 266 | MATa cyc1-512 ura3-52 trp2-1 HIS3::hisG tif4631::URA3 | tif4631Δ | *This Work |
| yBD 285 | MATa cyc1-512 *rrp4-1* ura3-52 trp2-1 *his3::hisG* | rrp4-1 | ySB 1065 Scott lab |
| yBD 298 | MATa cyc1-512 ura3-52 trp2-1 *HIS3::hisG HIS3-GAL::protA-RRP41* | HIS3-GAL10::protA-RRP41 | *This Work |
| yBD 300 | MATa cyc1-512 ura3-52 trp2-1 *his3::hisG upf1::URA3* | upf1Δ | *This Work |
| yBD 306 | MATa cyc1-512 ura3-52 trp2-1 *his3::hisGtrf4::URA3* | trf4Δ | *This Work |
| YBD334 | MATa cyc1-512 ura3-52 trp2-1 *his3::hisG* trf5::URA3 | trf5 Δ | *This Work |
| yBD454 | MATa cyc1-512 ura3-52 trp2-1 his3::hisG_YHR081w::kanMX4 (rrp47 Δ) | rrp47-Δ | *This Work |
| yBD455 | MATa cyc1-512 ura3-52 trp2-1 his3::hisG_YNR024w::kanMX4 (mpp6 Δ) | mpp6-Δ | *This Work |
| yBD458 | MATa cyc1-512 ura3-52 trp2-1 *his3::hisG air1*::URA3 | air1-Δ | *This Work |
| yBD161 | *MATa ade1 ade 2 lys2 gal1 ura3-52* | WT to 162 | Scott's Lab |
| yBD162 | MATa ade1 ade 2 lys2 gal1 ura3-52 rrp6:: URA3 | rrp6 Δ | Scott's Lab |
| yBD 163 | *MATa ade1 ade2 lys2 gal1 ura3-52 pap1-1* | pap1-1 | Scott's Lab |
| yBD 179 | *MATa ade1 ade2 lys2 gal1 ura3-52 pap1-1 rrp6::URA3* | pap1-1 rrp6Δ | Made By A.Maity |
| yBD117 | Mat a leu2-3,-112 ura3-52 his3-D 200 trp1-D901 lys2-801 | WT to 129 | Sarkar et. al 2018 |
| yBD 129 | Mata leu2-3,-112 ura3-52 his3-D 200 trp1-D901 lys2-801*rrp6::URA3* | rrp6-Δ | Sarkar et. al 2018 |
| yBD413 | Mata leu2-3,-112 ura3-52 his3-D 200 trp1-D901 lys2-801*rrp6::URA3* p*rrp6-3* | rrp6-3 | Sarkar et. al 2018 |
| yBD 65 | *MAT α ura3-1 ade2-1 his3-11,15 leu2-3,112 trp1-1* | WT to yBD66 | B-15142 |
| yBD 66 | *MAT α ura3-1 ade2-1 his3-11,15 leu2-3,112 trp1-1 dob1-1* | *MS157-15-dob1-1* | B-15143 |
| yBD 421 | MAT α ura3-1 ade2-1 his3-11,15 leu2-3,112 trp1-1 *rrp6*::URA3 | rrp6-Δ | *This work |
| yBD507 | MATa; ura3Δ0; leu2Δ0; his3Δ1; met15Δ0 RRP47 TAP:: KANMX4 | TAP-RRP47 | Gift from Rahmouni's Lab |
| yBD510 | MATa cyc1-512 ura3-52 trp2- 1 *his3*::hisG YHR081w::kanMX4 (rrp47Δ): rrp6Δ::URA3 (rrp6 Δ) | *rrp47Δ_rrp6Δ* | *This work |

***Table S2: List and Descriptions of Plasmids constructed and used***

| Plasmid no. | Description | Use | Reference |
|---|---|---|---|
| pBD19 | *pRS306-pBSKS URA3;amp$^R$* | | Sikorsky and Hieter (1989) |
| pBD 23 | *pRS 316 - pBSKS URA3; CEN6, ARS4; amp $^R$* | | Sikorsky and Hieter (1989) |
| pBD 24 | A 1.1 Kb *Hind*III fragment containing the transcribed region of URA3 gene in pUC19. | | This work[¥] |
| pBD 25 | A 3.0 kb SalI-BamHI fragment containing 1.1 kb URA3 gene inserted at HindIII site of CBC1 gene flanked by 5′ and 3′ flanking sequence of the same cloned in pUC 19. | CBC1 Disruptor | Das et al., 2000 |
| pBD 26 | A 3.0 kb fragment carrying 1.1 kb URA3 gene flanked by 5′ and 3′ flanking sequence of UPF1 gene in pBR322. | UPF1 Disruptor | Leeds et al .,1992 |
| pBD32 | A 5.5 kb fragment carrying 3.8 kb blaster sequence flanked by 5' and 3'flanking sequence of *CBC1* gene in pUC19. *CBC1* blaster. | CBC1-Blaster | Das et al., 2000 |
| pBD42 | A 4.1kb fragment containing the *CBC1* (from pBD 32) fragment is cloned in pCR 2.1 vector containing 5'and 3'part of *CBC1* ORF and Blaster. | CBC1-Blaster | Das et al., 2000 |
| pBD 45 | A 1.4 kb fragment carrying 1.1 kb URA3 gene flanked by 5′ and 3′ flanking sequence of RRP6 gene in pUC19. | RRP6 Disruptor | Briggs et al.,1998 |
| pBD 216 | A 3.8Kb blaster fragment from pBD 27(digested with BamHI, Bgl II) is cloned into pRS 313 (pBD 20) to get an HIS disruptor. Digestion with Bgl I & Kpn I gives the final HIS disruptor fragment. | HIS3 Disruptor | *This work |
| pBD225 | A 2.3 Kb PCR amplified (by OBD 407+ 408) Trf4 fragment from appropriate gene cloned in pJET1.2. | TRF4 in pJET1.2 | *This work |
| pBD226 | A 1.1 Kb URA3 fragment digested by *Hind*III from pBD24, inserted into *Eco*RI and *Eco*RV site of pBD225. | TRF4 Disruptor | *This work |
| pBD 248 | A 2.6 Kb Trf5 fragment (PCR Product by OBD 494+ 495) cloned in pJET1.2. Host DH5α | TRF5 in pJET1.2 | *This work |
| pBD 254 | 1.4 Kb URA3 fragment from SspI- NsiI site of pRS 306 is cloned into BstEII - NsiI site of pBD 248. PCR with oBD 494 - oBD 495 gives the 2.9 Kb TRF5 Disruptor fragment. | TRF5 Disruptor | *This work |
| pBD 266 | 1.4 Kb AIR1 fragment (by oBD 549+550) in pJET | AIR1 in pJET1.2 | *This work |
| pBD285 | A 1.6 Kb URA3 fragment from NsiI - XmnI restriction site of pRS 306 cloned into NsiI-MscI restriction site of pBD 266 gives AIR1 disruptor plasmid. | AIR1 Disruptor | *This work |

##### *Table S3. Oligonucleotides used for End Point PCR in this study*

| OBD NO. | OBD NAME | OBD SEQUENCE (5' to 3') | GENE TO AMPLIFY | REFERENCE |
|---|---|---|---|---|
| OBD 3 | BD-CBC1-D-S1 | 5'-ACTGTGTAAAGAAATGATGCCC-3' | CBC1 | Das et al.,2003 |
| OBD 4 | BD-CBC1-D-AS1 | 5' CGATATCCAATTCAATCTTCGC-3' | | |
| OBD 5 | BD-RRP6-D-S1 | 5' AGAATTTAGACAGGGG-3' | RRP6 | Briggs et al., 1998 |
| OBD 6 | BD-RRP6-D-AS1 | 5' CAT CGTCTCTTCTTGC-3' | | |
| OBD 19 | CBC1-BLASTER 5' JUNCTION-S1 | 5' CGAATGTAGTCCATCCTCCGAATC-3' | CBC1 BLASTER | Maity et al., 2016 |
| OBD 20 | CBC1-BLASTER 5' JUNCTION-AS1 | 5' TGTGCTCCTTCCTTCGTTCTTCCT-3' | | |
| OBD 25 | CBC1-BLASTER 3' JUNCTION-S2 | 5' TGATGATGACATTCCGGGTCTGGT-3' | CBC1 BLASTER | Maity et al., 2016 |
| OBD 24 | CBC1-BLASTER 3' JUNCTION-AS1 | 5' CATACCCAACTTTGACTACCTTGC-3' | | |
| OBD 43 | AM- RRP4-K-S1 | 5’-TGACGTATTCCTCTGTTGCTG-3’ | RRP4 | Maity et al., 2016 |
| OBD 44 | AM-RRP4-K-AS1 | 5’-AAAGAGGCACTACCATCTTGAAA-3’ | | |
| OBD 53 | SD TIF4631 ORF S2 | 5’-ATGATGGGCGCTATCCTCATCGC-3’ | TIF4631 | Das & Saha 2014 |
| OBD 54 | SD TIF4631 ORF AS2 | 5’-TATCGTAAACGAAGACTGGCGAAGC-3’ | | |
| OBD 79 | AM-Gal10-Rrp41-Ins-S2 | 5' TTCCCTGCGTCGCTTGCTGA 3' | RRP41 | Maity et al., 2016 |
| OBD 80 | AM-Gal10-Rrp41-Ins AS2 | 5’ACCGCCATCTTGCTCAAGGAC 3’ | | |
| OBD 254 | SD-UPF1-Forward | 5’-ATTTTAGTATCATCAGTTTCC-3’ | UPF1 | Das & Saha 2014 |
| OBD 255 | SD-UPF1-Reverse | 5’-CAACAAGTATAACTTGTTTCG-3’ | | |
| OBD 362 | AM_trf4_S1 . | 5'ACCTGTATTCATCCTCGGCTTCTTGTATATGATTCTAAAATAATGTGTGAAAAAAAAAATTTGATTCGGTAATCTCCGAGCAG-3' | TRF4 | Maity et al., 2016 |
| OBD 363 | AM_trf4_AS1 . | 5'ACACATTCTATCCAGGTACACAGTGATGTACAGTTCAGTGCATCATTTAAACAAAAAAGGCTAATTTGTGAGTTTAGTATACATGCATTTACTT-3' | | |
| OBD 405 | FORWARD:AC_air1_S1 | 5'GGCCAAGGACGGTATTTTGG-3' | AIR1 | *This work |
| OBD 406 | REVERSE:AC_air1_AS1 | 5'CACATGCAAGACGTAGACCCA-3' | | |
| OBD 407 | FORWARD:AC_trf4_S1 | 5'AAGTAAAGGATCCCAAGCGTGA-3' | TRF4 | Maity et al., 2016 |
| OBD 408 | REVERSE:AC_trf4_AS1 | 5'TTCGGCAAACGTGCCAATTT-3' | | |

| OBD NO. | OBD NAME | OBD SEQUENCE (5' to 3') | GENE TO AMPLIFY | REFERENCE |
|---|---|---|---|---|
| OBD 419 | AC_HIS_Del S1 | 5’-TTGGCCTCCTCTAGTACACTCT-3’ | HIS3 | *This work |
| OBD 420 | AC_HIS_Del AS1 | 5’-CAACCGCAAGAGCCTTGAAC-3’ | | |
| OBD 425 | AC_trf4 dsrptor_S2 | 5'-GTGCCGACATTTGAGGAGGA-3' | TRF4 | Maity et al., 2016 |
| OBD 426 | AC_trf4 dsrptor_AS2 | 5'-CGCTAAGGGCAGCGATTCTA-3' | | |
| OBD 429 | AC_trf4 dsrptor_S1 | 5’-AGTCTGCACAAGACCCTTCG-3’ | TRF5 | Maity et al., 2016 |
| OBD 430 | AC_trf4 dsrptor_AS1 | 5’-ACGTTACGATAGAGCCAGCG-3’ | | |
| OBD 494 | AM-TRF5-S1 | 5’-GCTAACACCACCACCCATGA-3’ | TRF5 | *This work |
| OBD 495 | AM-TRF5-AS1 | 5’-GGGAAAATACCCACGCGTTT-3’ | | |

***Table S4: Oligonucleotides used for RT PCR in this study***

| OBD NO. | OBD NAME | OBD SEQUENCE (5' to 3') | GENE TO AMPLIFY | REFERENCE |
|---|---|---|---|---|
| OBD 168 | AM -SCR1-S2 | 5’ TTGTGGCAACCGTCTTTCCT 3’ | SCR1 | *This work |
| OBD 169 | AM-SCR1-AS2 | 5’ CCGAAGCGATCAACTTGCAC 3’ | | |
| OBD 268 | US-ACT1-RT-S1 | 5’-GCCGAAAGAATGCAAAAGGA-3’ | ACT1 | *This work |
| OBD 269 | US-ACT1-RT-AS1 | 5’-TCTGGAGGAGCAATGATCTTGA-3’ | | |
| OBD 343 | AM-RT-5s_S1 | 5'-GGTTGCGGCCATATCTACCA-3' | MATURE 5S | *This work |
| OBD 344 | AM-RT-5s_AS1 | 5'-ACTACTCGGTCAGGCTCTTACCA-3' | | |
| OBD 188 | AM-5.8S-RT-S1- | 5’-AAC AAC GGA TCT CTT GGT TCT-3’ | MATURE 5.8S | *This work |
| OBD 189 | AM-5.8S-RT-AS1 | 5’-AAA TGA CGC TCA AAC AGG CA-3’ | | |
| OBD 341 | AM-RT-18s_S2 | 5'-GATCGGGTGGTGTTTTTTTAATG-3' | MATURE 18S | *This work |
| OBD 342 | AM-RT-18s_AS2 | 5'-CTCCCCCCAGAACCCAAA-3' | | |
| OBD 553 | AC_25s_RT_S1 | 5’-GGACTGAGGACTGCGACGTAA -3’ | MATURE 25S | *This work |
| OBD 554 | AC_25s_RT_AS1 | 5’- TCAAGACGGGCGGCATAT -3’ | | |
| OBD 578 | AC_snRNA U1_ S1 | 5'-ACGGCAGATTCGAATGAACTTAA-3' | snRNA U1 | *This work |
| OBD 579 | AC_snRNA U1_ AS1 | 5'-TACAATCCCGACCAAATAATCTCA-3' | | |
| OBD 580 | AC_snRNA U4_ S1 | 5'-AGGATTCGTCCGAGATTGTGTT-3' | snRNA U4 | *This work |
| OBD 581 | AC_snRNA U4_ AS1 | 5'-CATGAGGAGACGGTCTGGTTTAT-3' | | |
| OBD 582 | AC_SNR 10_S1 | 5'-CGATCTTGGGTGCAACAGTCT-3' | SNR 10 | *This work |
| OBD 583 | AC_SNR 10_AS1 | 5'-TCATCCGGGCACACGAA-3' | | |
| OBD 584 | AC_SNR 13_S1 | 5'-TGAGTGCATTTGGCTCGAGTT-3' | SNR 13 | *This work |
| OBD 585 | AC_SNR 13_AS1 | 5'-GCTTGAGTTTTTCCACACCGTTA-3' | | |

| OBD NO. | OBD NAME | OBD SEQUENCE (5' to 3') | GENE TO AMPLIFY | REFERENCE |
|---|---|---|---|---|
| OBD 600 | AC_snRNA U6_S1 | 5’-TCGTGGACATTTGGTCAATTTGA-3’ | snRNA U6 | *This work |
| OBD 601 | AC_snRNA U6_AS1 | 5’-TTTGTAAAACGGTTCATCCTTATGCA-3’ | | |
| OBD 638 | AC_pre 5s_S1 | 5'-GGAAACGGTGCTTTCTGGTAGA-3' | PRE 5S | *This work |
| OBD 639 | AC_pre 5s_AS1 | 5'-ATCACCTGCGTTTCCGTTAAA-3' | | |
| OBD 640 | AC_pre 18s_S1 | 5'-AGTCGTAACAAGGTTTCCGTAGGT-3' | PRE 18S | *This work |
| OBD 641 | AC_pre 18s_AS1 | 5'-CTTGCCAAAACAAAAAAATCCAT-3' | | |
| OBD 642 | AC_pre 5.8s_S1 | 5'-CATCGAATCTTTGAACGCACAT-3' | PRE5.8S | *This work |
| OBD 643 | AC_pre 5.8s_AS1 | 5'-GGCCAGCAATTTCAAGTTAACTC-3' | | |
| OBD 644 | AC_pre 5.8s Margin_S1 | 5'-CATCGAATCTTTGAACGCACAT-3' | PRE 5.8S II | *This work |
| OBD 645 | AC_pre 5.8s Margin_AS1 | 5'-AGAAGGAAATGACGCTCAAACAG-3' | | |
| OBD 646 | AC_pre 25s_S1 | 5'-CCTTGTTGTTACGATCTGCTGAGA-3' | PRE 25S | *This work |
| OBD 647 | AC_pre 25s_AS1 | 5'-TGCCAGTACCCACTTAGAAAGAAATAA-3' | | |
| OBD 658 | AC-ITS1-5.8s-S1 | 5'-CTGTGGAGTTTTCATATCTTTGCAA-3' | ITS1 | *This work |
| OBD 659 | AC-ITS1-5.8s-AS1 | 5'-TACCTCTGGGCCCCGATT-3' | | |
| OBD 677 | AC_RRP6_DEL_F1_ | 5'-TTATGTAAAACAAGCGTATTTTTTTATTTAT-3' | RRP6 | *This work |
| OBD 678 | AC_RRP6_DEL_R1 | 5'-TGGAATGGGAGTAGCTTCCT-3' | | |
| OBD 679 | AC_RRP6_DEL_R2 | 5'-TGGAATGGGAGTAGCTTC-3' | | |
| OBD 680 | AC pre U1 Forward | AACGGGTGGATCTTATAATTTTTGA | snRNA U1 | *This work |
| OBD 681 | AC pre U1 Reverse I | AGAAGCATGAAACTTTAAAAGTTTCAGTAC | | |
| OBD 682 | AC pre U1 Reverse II | AAACACATCACCAGATATAGGACTGAA | | |
| OBD 683 | AC_ pre U1 Reverse III | TCTCAATGTAGCGTATAACAGGTTTCA | | |
| OBD 684 | AC_pre U4 F | TCGGTGTTCGCTTTTGAATACTT | snRNA U4 | *This work |
| OBD 685 | AC_pre U4 R I | CCATTTTAGTTGCCATTTCTGTATTACTT | | |
| OBD 686 | AC_pre U4 R II | GTGAACTCACATTAGAGAATTCGG | | |
| OBD 687 | AC_pre U4 R III | CAATTCCAGCTTCTATAACGTGAAC | | |
| OBD 688 | AC_ pre U6 Forward | CAGTTCCCCTGCATAAGGATGA | snRNA U6 | *This work |
| OBD 689 | AC_ pre U6 Reverse | GTCACGATACTTCACTCGATGATAAAAA | | |
| OBD 690 | AC_ pre SNR13 Forward | GAGTTGCTGTTTGGCTTTTGC | PRE snR 13 | *This work |
| OBD 691 | AC_ pre SNR13 Reverse | ATTTTCTACGGGGAAGTTAAAAGGTC | | |
| OBD 692 | AC_ pre SNR13 Reverse II | AGCGCTACGATACAATGTAAGAAGG | | |
| OBD 698 | AC_RRP47 TAP Forward: | TAAGGAAAGCAAGTAGTAAGAAAAGTAAAAG ATTGGATAAAGTTGGAAAAAAGAAAGGAGGG AAGAAGTCATCCATGGAAAAGAGAAG | RRP47 | *This work |
| OBD 699 | AC_RRP47 TAP Reverse: | ATTTTTGCATTTGTGCTCTCACATCACCTTTAA TCATTTTTTCACTCATGTACCAGTATACGTCGA CCTTACGACTCACTATAGG | | |

| OBD NO. | OBD NAME | OBD SEQUENCE (5' to 3') | GENE TO AMPLIFY | REFERENCE |
|---|---|---|---|---|
| OBD 700 | AC_RRP6 TAP_Forward: | TAGTAATGGACCAAGGGCAGCTAAAAAGAGG AGGCCTGCCGCCAAAGGTAAGAATCTGTCATT TAAAAGGTCATCCATGGAAAAGAGAAG | RRP6 | *This work |
| OBD 701 | AC_RRP6 TAP_Reverse: | GATGAATTTAGAGGTCTTAAATGAAAATTACC ATAATTTATAAATAAAAAAATACGCTTGTTTTA CATAATACGACTCACTATAGGG | | |
| OBD 702 | AC_5s R1: | GGCTCTTACCAGCTTAACTACAGTTGA | MATURE 5S | *This work |
| OBD 703 | AC_5s F2: | GGTAAGAGCCTGACCGAGTAGTG | | |
| OBD 704 | AC_5s R2: | AAAGATTGCAGCACCTGAGTTTC | | |
| OBD 705 | AC_pre 5s F2: | GGCTTCCTATGCTAAATCCCATAAC | PRECURSOR 5S | *This work |
| OBD 706 | AC_pre 5s R2: | GCAGCTGGATAGTGCGAATTTT | | |
| OBD 707 | AC_5.8s R2: | GGCGCAATGTGCGTTCA | MATURE 5.8S | *This work |
| OBD 708 | AC_5.8s F3: | CATCGAATCTTTGAACGCACAT | | |
| OBD 709 | AC_5.8s F4 MARGIN: | CGTCATTTCCTTCTCAAACATTCTG | | |
| OBD 710 | AC_Pre 5.8s R4: | GGCCAGCAATTTCAAGTTAACTC | PRECURSOR 5.8S | *This work |
| OBD 711 | AC_Pre 5.8s F5: | GGCCTTTTCATTGGATGTTTTT | | |
| OBD 712 | AC_Pre 5.8s R5: | AAACGACCGTACTTGCATTATACCT | | |
| OBD 713 | AC_Pre 5.8s F6: | CTGCGGCTAATCTTTTTTTATACTGA | | |
| OBD 714 | AC_Pre 5.8s R6: | GTTCGCCTAGACGCTCTCTTCTTA | | |
| OBD 715 | AC_U1 F2: | TCCTTGGTCACACACACATACG | snRNA U1 | *This work |
| OBD 716 | AC_U1 R2: | GGGAATGGAAACGTCAGCAA | | |
| OBD 717 | AC_Pre U1 F III: | TGCTTCTATTTTCTTCATTTCAGTCCTA | PRE snRNA U1 | *This work |
| OBD 718 | AC_Pre U1 F4: | TACGCTACATTGAGACAAGACATTGT | | |
| OBD 719 | AC_Pre U1 R4: | TGCGGCTCTCTTAAGCACATT | | |
| OBD 720 | AC_Pre U4 F2: | TTAGGGATACCGGACTGAAACATT | PRE snRNA U4 | *This work |
| OBD 721 | AC_Pre U4 F3: | CGAATTCTCTAATGTGAGTTCACGTT | | |
| OBD 722 | AC_Pre U4 F4: | TGTTCTCCTTCTTGTGTGTCTTTCTT | | |
| OBD 723 | AC_Pre U6 F2: | ACCCAGCGTACAGCAGTGTATCT | PRE snRNA U6 | *This work |
| OBD 724 | AC_Pre U6 R2: | TCGAACGCGAGACAATTTTCTA | | |
| OBD 725 | AC_snR10 F2: | GCCATTCGTAACACGTACAGTATCTC | MATURE snR 10 | *This work |
| OBD 726 | AC_snR10 R2: | CCACATTCTTCATGGGTCAAGA | | |
| OBD 727 | AC_Pre snR10 F1: | TGTATATATGAGTGGCTTGGAATGC | PRE snR 10 | *This work |
| OBD 728 | AC_Pre snR10 R1: | AAATTGCGTCAACCGAAGGA | | |
| OBD 729 | AC_Pre snR13 F2: | TTCCCCGTAGAAAATCTTAGTAATCC | PRE snR 13 | *This work |
| OBD 730 | AC_Pre snR13 R3: | GCCAAACCCAACGTACTAACATC | | |
| OBD 731 | AC_Pre snR13 F3: | GCGCTGCATATATAATGCGTAAAAT | | |

| OBD NO. | OBD NAME | OBD SEQUENCE (5' to 3') | GENE TO AMPLIFY | REFERENCE |
|---|---|---|---|---|
| OBD 732 | AC_U2 F1: | TGGCACCCAAAATAATAAAATGG | snRNA U2 | *This work |
| OBD 733 | AC_U2 R1: | TGAGACCTGACATTAGCGGAAA | | |
| OBD 734 | AC_U2 F2: | CCCCAAGTATCGGCCAAAGT | snRNA U2 | *This work |
| OBD 735 | AC_U2 R2: | ACCCTACACCCCCTCAAACC | | |
| OBD 736 | AC_U2 F3: | CCGGCGGCATCAAGAA | snRNA U2 | *This work |
| OBD 737 | AC_U2 R3: | AGGGTCGCGACGTCTCTAACT | | |
| OBD 738 | AC_Pre U2 F4: | TTTTTCCTTTGACTTCGCATGA | PRE snRNA U2 | *This work |
| OBD 739 | AC_Pre U2 R4 | TTTGGTTGCGTGGTATATAGAATCTC | | |
| OBD 740 | AC_Pre U2 F5 | TGGCGGCATACTCTTTCTTGA | PRE snRNA U2 | *This work |
| OBD 741 | AC_Pre U2 R5 | AGGCAATGGGAAGCAGCTAA | | |
| OBD 742 | AC_U5 F1 | GGGAGGTCAACATCAAGAACTGT | snRNA U5 | *This work |
| OBD 743 | AC_U5 R1 | GATGGTTCTGGTAAAAGGCAAGA | | |
| OBD 744 | AC_U5 F2 | TCCGGGTGTTGTCTCCATAGA | snRNA U5 | *This work |
| OBD 745 | AC_U5 R2 | AGGGCAGAAAAGTTCCAAAAAAT | | |
| OBD 746 | AC_Pre U5 F3 | GGAGGGCGTTTATCTTTTCTATTTTATT | PRE snRNA U5 | *This work |
| OBD 747 | AC_Pre U5 R3 | CATGGACTCATGAATCAAATTTGTAGA | | |
| OBD 748 | AC_Pre U5 F4 | GAGTCCATGGAACAAATATATAGAACTCA | PRE snRNA U5 | *This work |
| OBD 749 | AC_Pre U5 R4 | GCAGTCAGGATAAAAGCAAATGC | | |

***Table S5: Different Primary antibody used for the work and their respective secondary Antibody***

| Sl. No. | Primary Antibody | 1º Ab Dilution | Secondary Antibody | 2º Ab Dilution |
|---|---|---|---|---|
| 1 | Anti-Rrp6 | 1:1000 | Anti- Rabbit | 1:3000 |
| 2 | Anti-Tap (for Rrp47) | 1:1000 | | |
| 3 | Anti-Rrp4 | 1:1500 | | |
| 4 | Anti- Alpha Tubulin | 1:2000 | Anti-Mouse | 1:3000 |

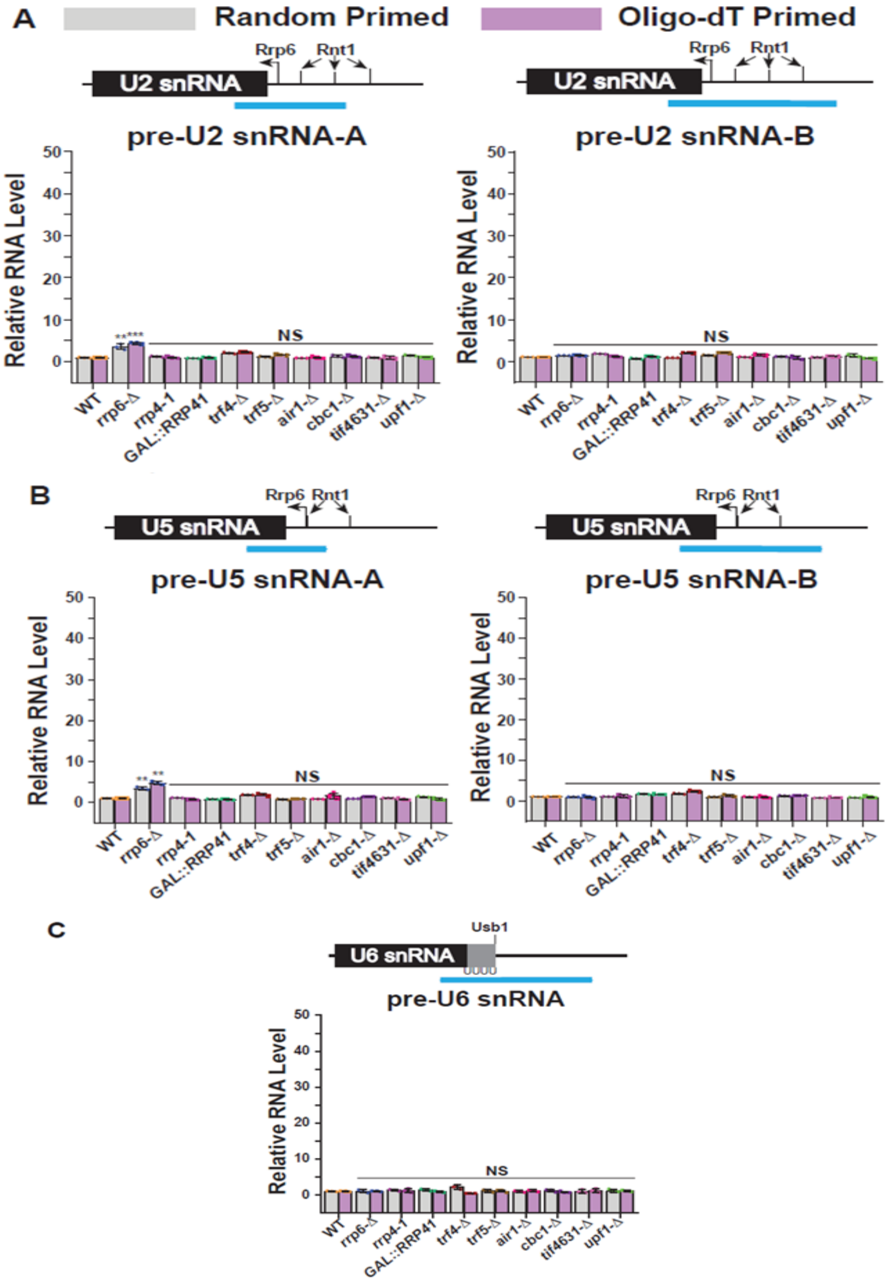


**Supplementary Figure S1: Steady-state levels of 3′-extended precursor forms of different test snRNAs in various yeast strains.** Scattered/Bar plot revealing the steady-state levels of 3′-extended precursor forms of (A) snRNA U2, (B) snRNA U5 and (C) snRNA U6 in the indicated yeast strains. The RNA levels were estimated from the 2 ng cDNA samples prepared using random hexanucleotide primers (grey bars) or oligo-$dT_{30}$ anchor primer (pink bars). Various amplicons used in subsequent qRT-PCR reactions covering partly the mature region and partly the 3′-extended precursor regions of each RNA species are shown in the schematic figures presented on top of each graph. Different 3′-end processing sites indicating trimming by Rrp6p and cleavage by Rnt1p and RNaseIII are denoted in the respective schematic figures. *SCR1* (in case of Random Primer) and *ACT1* mRNA (in case of Oligo dT Primer) were used as the internal control. Abundance of these ncRNAs in *upf1*-Δ yeast strain was used as a negative control. Normalized values of each of the ncRNAs in the wild type yeast strain were set to 1. Three independent cDNA preparations (biological replicates, n=3) were used to determine the levels of various ncRNAs. The statistical significance of difference reflected in the ranges of P values are presented with the following symbols, *<0.05, **<0.005, and ***<0.001; NS, not significant.

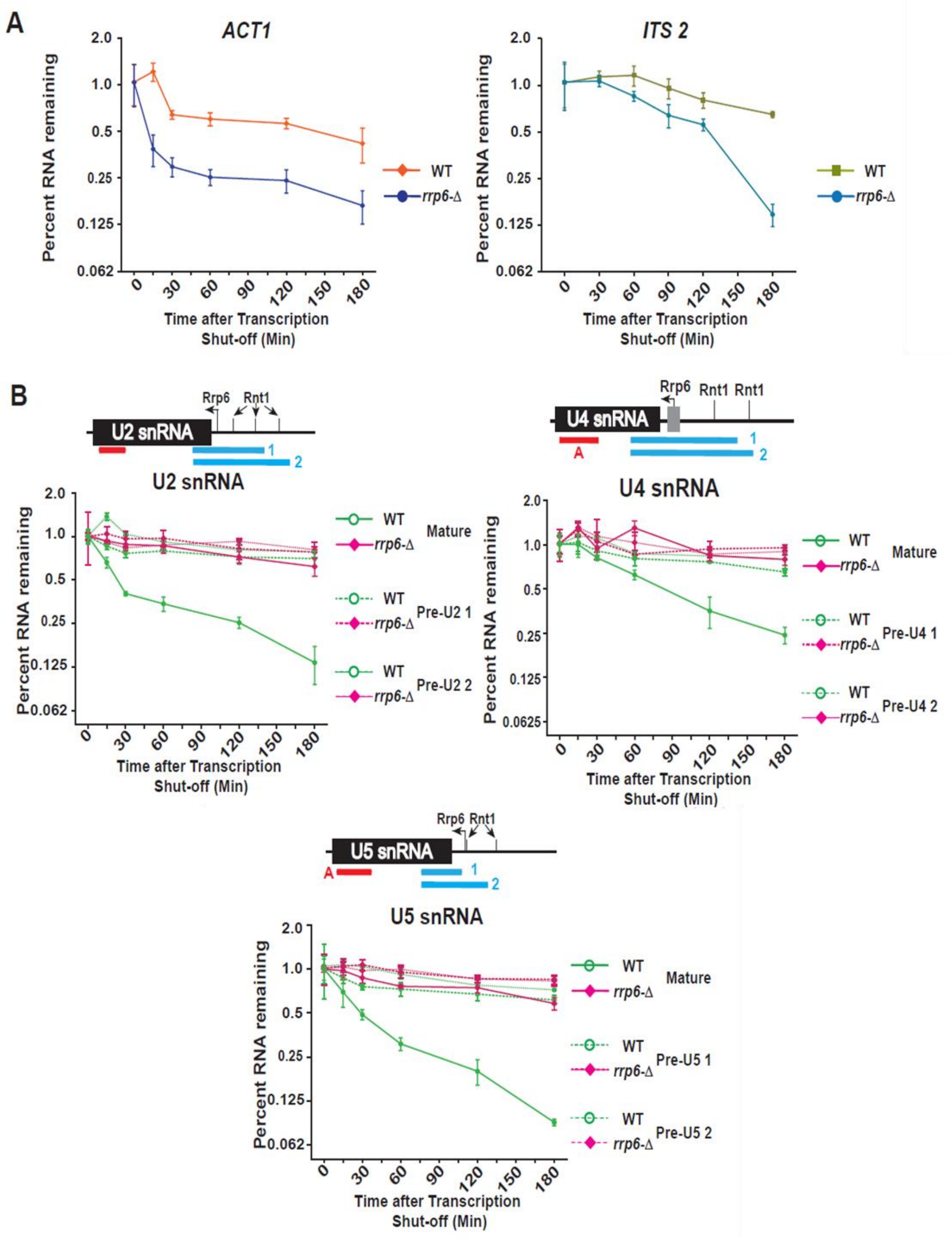


**Supplementary Figure S2: Transcription Shut-off experiment showing polyadenylated mature forms of the small non-coding RNAs undergo an active degradation by Rrp6p.** Time kinetics of the steady-state levels of total and precursor small non-coding RNAs following the transcription shut-off revealing their decay rates in wild-type and rrp6-Δ strains. The decay rates were determined from four independent experiments (biological replicates, n=4) by qRT-PCR analysis (using amplicons indicated in thick red/blue/green lines below each gene sequence) and the signals were normalized to either SCR1 RNA (in case of inhibition of RNA Pol-II) or ACT1 mRNA (in case of inhibition of RNA Pol-I and III). Normalized signals (mean values ± SD) were presented as the fraction of remaining RNA (with respect to normalized signals at 0 min) as a function of time of incubation in the presence of either transcription inhibitors BMH-21 (for RNA Pol I Transcripts); 1,10-phenanthroline (for RNA Pol II Transcripts); and ML-60218 (for RNA Pol III Transcripts). WT and rrp6-Δ are denoted with violet/blue, green/pink, and black/red colored lines for rRNA, snRNA, and snoRNA respectively. Time kinetics of the steady-state levels of ACT1 and ITS1 are taken as control for RNA Pol II and Pol I decay respectively. RNAs qRT-PCR assays, Total RNA/cDNA isolations were carried out as described in materials and methods.

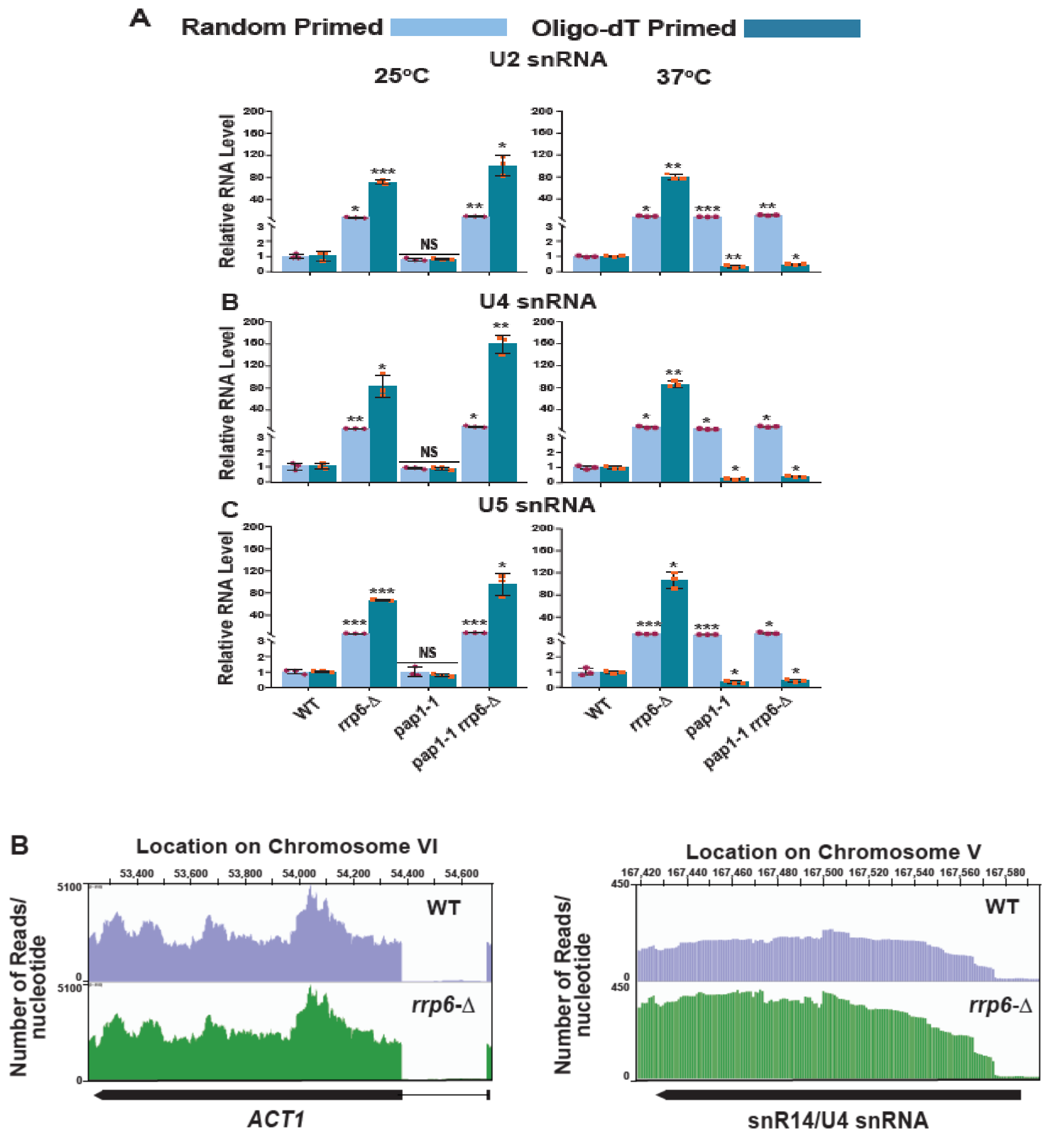


**Supplementary Figure S3: The canonical Poly(A) polymerase Pap1p play vital role in the polyadenylation of the smaller ncRNAs.** (A) Scattered/Bar plot revealing the steady-state levels of various low molecular weight ncRNAs estimated from the 2 ng cDNA samples prepared using random hexanucleotide primers (sky blue bars) or oligo-dT$_{30}$ anchor primer (indigo blue bars) by qRT-PCR from WT and yeast strains carrying mutations in the *RRP6* (*rrp6*-Δ), *PAP1* (*pap1-1*), *pap1-1 rrp6*-Δ double mutant alleles. The strains were pre-grown at 25ºC followed by splitting the culture into two halves. One half was continued to grow at 25°C for 12 hours and a 12h shift to 37ºC were performed to the other half of the culture before harvesting them. Total RNA, cDNA isolation from them followed by qRT-PCR reaction are carried out as described in materials and methods. *SCR1* (in case of Random Primer) and *ACT1* mRNA (in case of Oligo dT Primer) were used as the internal loading control. Normalized values of each of the ncRNAs in the wild type yeast strain were set to 1. Three independent cDNA preparations (biological replicates, n = 3) were used to determine the levels of various ncRNAs. The statistical significance of difference reflected in the ranges of P values estimated from Student's two-tailed t tests for a given pair of test strains for every message are presented with the following symbols, *<0.05, **<0.005, and ***<0.001; NS, not significant. (B) Analysis of previously done RNA-seq data (Accession Number GSE135056) revealed a dramatic accumulation of sense strand reads corresponding to the mature region of several small nucleolar RNAs. Graphical representation showing the relative amount of sense strand reads mapped to the genomic locus corresponding to ACT1 and snR14/snRNA U4. The location of transcripts and the direction of transcription are shown below the graph (drawn in scale) by the solid black arrow-headed rectangles.
